# Microbial profiling of Drosophila in French Guiana reveals candidate microbial taxa associated with native and invasive host

**DOI:** 10.64898/2026.06.12.731834

**Authors:** Thibault T. Laffargue, Nicolas Pollet, Louise Brousseau, Wolfgang J. Miller, Aurélie Hua-Van, Mathieu Chouteau

**Affiliations:** Laboratoire Écologie, Évolution, Interactions des systèmes Amazoniens, University of French Guiana, CNRS, IFREMER, Cayenne, France; EGCE, University Paris-Saclay, CNRS, IRD, Gif-sur-Yvette, France; UMR AMAP, University of Montpellier, IRD, CIRAD, CNRS, INRAE, Montpellier, France; Center of Anatomy and Cell Biology, Medical University of Vienna, Austria

**Author notes:** Corresponding authors: Mathieu Chouteau, Laboratoire Écologie, Évolution, Interactions des systèmes Amazoniens, University of French Guiana, CNRS, IFREMER, Cayenne, France;., Aurélie Hua-Van, EGCE, University Paris-Saclay, CNRS, IRD, Gif-sur-Yvette, France;. These authors contributed equally as senior authors (Aurélie Hua-Van, Mathieu Chouteau).

**Keywords:** microbiome, Drosophila, Neotropical, host adaptability, invasive alien species, French Guiana

## Abstract

Invasive alien species (IAS) represent a major threat to biodiversity, human health, and economy. Yet how host-associated microbiomes differ between invasive and native species remains poorly understood in natural systems. Here, we investigated the bacterial and fungal microbiomes of IAS and native *Drosophila* species collected along an anthropisation gradient in French Guiana. Analyses of *Drosophila* communities revealed that a strong decline in IAS occurrence from coastal anthropised habitats toward rainforest localities. Using bacterial 16S rRNA and fungal ITS metabarcoding of pooled flies, we assessed microbial diversity, composition, and structure across host species, host category (invasive vs. native), and locality. Bacterial communities associated with IAS harboured reduced alpha diversity and differed in community structure from those associated with native Drosophila, while bacteriome variation was also associated with host phylogenetic grouping and locality. Because host category was partially confounded with host phylogeny and locality, these differences cannot be attributed uniquely to invasion status. In contrast, fungal communities were primarily associated with locality and host species. We further identified core and exclusive taxa, compared bacteriome composition between native and IAS hosts using global reference datasets, and integrated information from the microbiome literature. This approach identified bacterial and fungal candidate taxa that were preferentially or exclusively associated with invasive or native hosts. Overall, our results characterise microbiome variation between invasive and native Drosophila in French Guiana and identify candidate microbial taxa for future functional investigation.

## Introduction

Animals and plants introduced outside their natural range that establish and spread are termed invasive alien species (IAS), sometimes defined to include their economic, environmental, and health impacts (IUCN, 2000; Richardson et al., 2000; Occhipinti-Ambrogi & Galil, 2004; Pyšek et al., 2004; Blackburn et al., 2011). IAS pose a major threat to biodiversity by reducing species richness and native abundance, increasing extinction risk, and disrupting trophic networks, while also altering the genetic composition of native populations through hybridisation, introgression, and selective pressures (Sakai et al., 2020; Biedrzycka et al., 2012; Todesco et al., 2016; Pyšek et al., 2020). They also threaten human and animal health by acting as vectors of pathogens and parasites, either introducing them into invaded areas or amplifying those already present (Chinchio et al., 2020). Ultimately, IAS can also impact the economy by destroying agricultural resources and generating substantial management costs (Haubrock et al., 2021; Roiz et al., 2024; Bradshaw et al., 2024; Bacher et al., 2025; Monguilod and Gallardo, 2026).

Understanding IAS biology is therefore a priority. In particular, invasion success has been linked to shared life-history traits such as short life cycles and high phenotypic plasticity (Davidson et al., 2011; Capellini et al., 2015; Allen et al., 2017). In parallel, other studies emphasise the importance of propagule pressure, including the number and frequency of introductions (Lockwood et al., 2005; Cassey et al., 2018). Another widely discussed explanation is the “enemy release hypothesis” which attributes invasion success to the absence of natural predators in the invaded area (Jeffries & Lawton, 1984; Roy et al., 2011). In addition to these well-established mechanisms, increasing evidence suggests that the microbiome may also play a role in the establishment success of IAS (Kelly et al., 2009; Taraschewski et al., 2006; Poulin et al., 2011; Hatcher et al., 2012; Lymbery et al., 2014; Vilcinskas et al., 2015; Lu et al., 2016; Le Roux et al., 2017; Martignoni et al., 2024; Romeo et al., 2025; Martignoni et al., 2025).

In the context of biological invasions, introduced species are generally accompanied by part of their microbiome, a phenomenon known as co-introduction (Dawkins, 1982; Lu et al., 2016; Le Roux et al., 2017). In addition, microorganisms from the invaded environment, or exchanged with native species, may be incorporated into the IAS microbiome (Taraschewski et al., 2006; Kelly et al., 2009; Poulin et al., 2011; Hatcher et al., 2012; Lymbery et al., 2014; Vilcinskas et al., 2015; Lu et al., 2016; Martignoni et al., 2024). The microbiome is therefore often considered to confer an extended phenotype on the host and to contribute to its evolutionary history and environmental adaptation (Calderon-Cortés et al., 2012; Davenport et al., 2017; Kolodny et al., 2020; Henry et al., 2021; Petersen et al., 2023; Bordenstein et al., 2024). As part of this extended phenotype, the microbiome in insects aids digestion, protects against parasites and pathogens, mediates inter- and intraspecific communication, influences mate finding and reproductive success, and provides essential amino acids as well as metabolic compounds and nutrients (Douglas et al., 2001; Douglas, 2011; Engel and Moran, 2013; Douglas, 2015; Gupta et al., 2020; Barron et al., 2023). Consequently, changes in the composition of co-introduced or locally acquired microorganisms could modify host-associated functions and potentially influence traits relevant to establishment and persistence. However, associations between microbiome variation and biological invasions remain insufficiently investigated in field settings, particularly across native and invasive hosts occurring in the same region.

To address this gap, the genus *Drosophila* provides a particularly powerful model for investigating host-associated microbiome variation in an invasion context. This genus, which comprises approximately 2,000 species of fruit flies, and in particular *D. melanogaster*, has been a universal model organism for over a century owing to its short life cycle, ease of laboratory rearing, and genetic tractability (Dobzhansky, 1937; Roberts, 2006). For decades, *Drosophila* has served as a model for microbiome studies, first under laboratory conditions and more recently in natural environments (Chamilos et al., 2006; Cox and Gilmore, 2007; Chandler et al., 2011, 2012; Broderick and Lemaitre, 2012; Douglas et al., 2018; Brown et al., 2023; Comeault et al., 2024; Cho and Kang, 2025). In the wild, the *Drosophila* microbiome is typically dominated by *Acetobacteraceae*, *Lactobacillaceae*, *Enterobacteriaceae*, *Enterococcaceae*, *Pseudomonadaceae*, and environmental yeasts (Staubach et al., 2013; Brown et al., 2023; Ludington et al., 2025; Bhandari et al., 2025; Medeiros et al., 2025). Given this extensive body of knowledge, *Drosophila* represents an appropriate system for for comparing microbiome associations between native and invasive hosts.

Within this framework, South America is a particularly relevant system due to its high diversity of both invasive alien and Neotropical *Drosophila* species (Dobzhansky and Pavan, 1943; Val and Sene, 1980; Brncic, 1987; Caracristi, 2003; Gottschalk, 2008; Tidon and Almeida, 2016). More specifically, in French Guiana, on the northeastern coast of South America between Brazil and Suriname, preliminary captures conducted in 2019 identified several IAS and native species (Miller and Hua-Van, unpublished; Laffargue et al., in preparation). The coexistence of IAS and native species, combined with the tractability of the system, provides an opportunity to compare their associated microbiomes within the same Neotropical region. Nevertheless, despite the large number of studies on *Drosophila* microbiomes, Neotropical systems remain comparatively understudied.

In this study, we investigated the composition and structure of bacterial and fungal communities associated with invasive and native Neotropical *Drosophila* species in French Guiana. We first examined how *Drosophila* communities are distributed along an anthropisation gradient, given the known association between human activity and IAS occurrence. We then explored variation in microbiome alpha diversity, composition, and community structure across localities, host species, and host category (IAS vs. native). Finally, we identified core and exclusive microbial taxa contributing to differences between IAS and native hosts, thereby generating a set of candidate taxa for future functional investigation.

## Materials and methods

### Collection of individuals

To assess fruit fly diversity, traps were deployed for 48 h in four localities along an anthropisation gradient in French Guiana: Cayenne (100 traps, 10 sessions), Kaw (140 traps, 3 sessions), the Bélizon road (68 traps, 2 sessions), and the Nouragues (248 traps, 5 sessions) (**Figure 1A;** APA NOR: TREL2400559S/814). Cayenne, the largest city and main entry point for goods and human activity, is characterised by a high level of anthropogenic disturbance (coordinates: 4.94074, -52.32433; 4.93094, -52.29725). Kaw and the Bélizon road are more remote forested sites that remain connected to Cayenne by road. At Kaw, traps were placed 1– 1.5 km from the village (4.47756, -52.03438 to 4.47276, -52.03803), while at Bélizon they were deployed along transects perpendicular to the road. The Nouragues reserve represents a remote and minimally disturbed site, with traps positioned at least 30 minutes’ walking distance from the CNRS Inselberg station, located tens of kilometres from the nearest human settlement (4.07883, -52.68716 to 4.07496, -52.69247).

**Figure 1.**
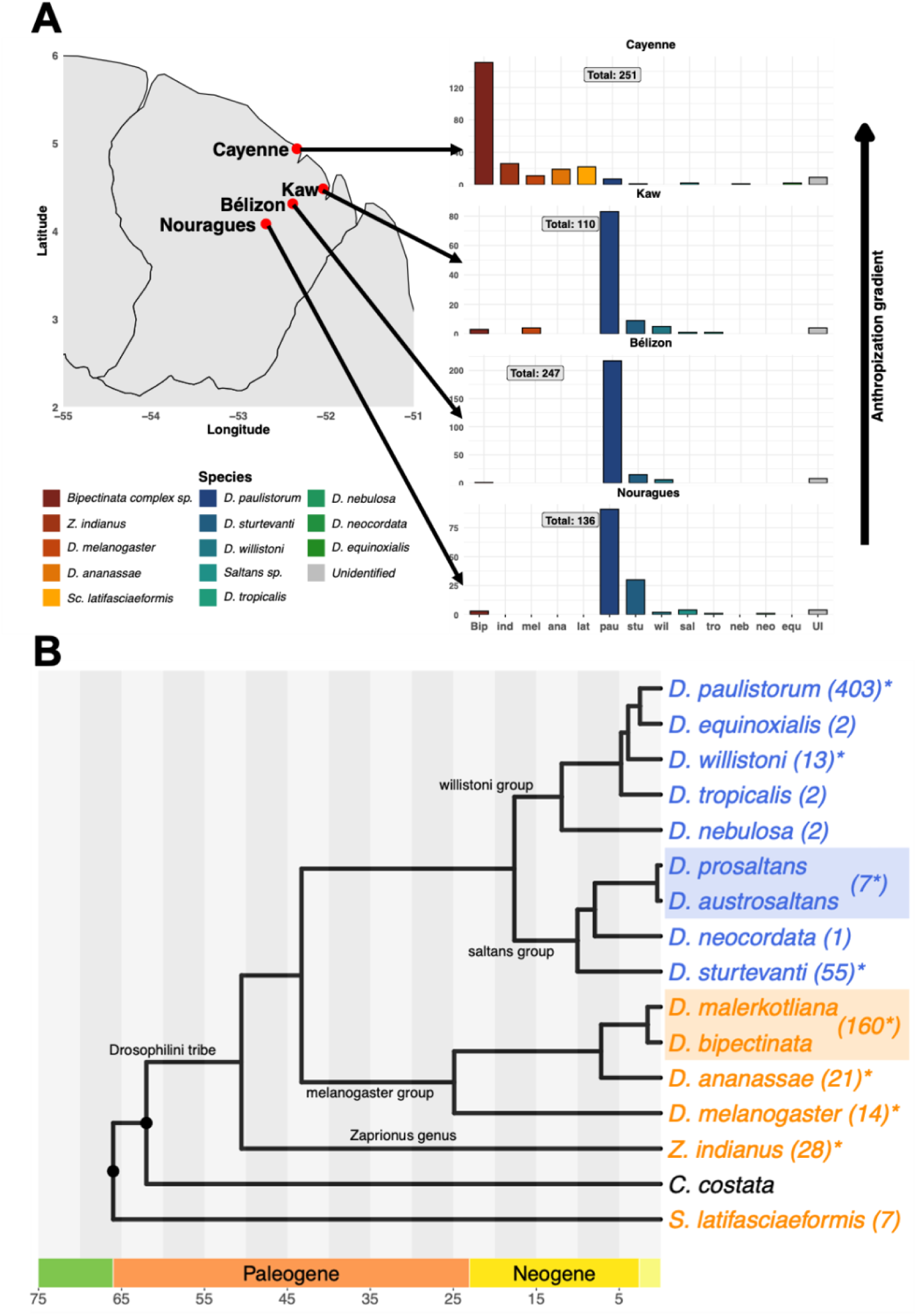
French Guiana *Drosophila* community composition and phylogeny. (A) *Drosophila* community composition at each locality. Invasive alien species are in red-orange, native species in blue-green, and unidentified in grey. Each bar chart corresponds to a single locality, and the total number of individuals collected is indicated in the center of the chart. (B) Phylogenetic relationships of captured *Drosophilidae* species. Invasive species are in orange and native neotropical species in blue. *Chymomyza costata* was added for calibration purposes. The number of captured specimens is indicated into parentheses. Note that some specimens cannot be distinguished based on the *COI* sequences and are referred to as the saltans subgroup (*D. autrosaltans/D. prosaltans*) or the bipectinata subcomplex (*D. malerkotliana/D. bipectinata*). * Denotes species/groups that were further considered for microbiome analysis. See methods for details about tree reconstruction.

All traps consisted of plastic bottles constructed following a standard protocol, with the addition of rain protection and a cotton barrier to limit potential contamination of the fly microbiome by the bait (Reed, 1938; Williams & Miller, 1952). Two types of fruit were used as bait: pomelo and pineapple.

Traps were deployed between May 22 and August 29, 2023, and retrieved after 48 h, with sampling alternating among localities. Due to logistical constraints, all sampling at the Nouragues was conducted in August. All flies captured within a trap were collected using an insect aspirator and transferred together into a single empty *Drosophila* vial. In the laboratory, samples were briefly frozen (10 min at −20 °C) to immobilise individuals, which were then sexed, morphologically identified to species group, and preserved individually in 0.2 mL tubes containing ZymoResearch DNA/RNA Shield at −20 °C.

### Taxonomic identification of collected flies

Because morphological identification alone did not provide reliable species-level resolution, flies were individually characterised by sequencing a *COI* DNA fragment to infer species identity. Flies were removed from DNA/RNA Shield, rinsed with sterile water, and three to four legs were collected from each individual into sterile 96-well PCR plates. The remaining body was immediately returned to DNA/RNA Shield for microbiome analyses.

For each sample, 20 µL of squishing buffer (H₂O, 0.01 M Tris-HCl pH 8, 1 mM EDTA, 25 mM NaCl) supplemented with 1% proteinase K was added, tissues were homogenized using a pipette tip and incubated overnight at 37 °C, followed by a 2 min incubation at 95 °C to inactivate proteinase K. PCR amplification was performed using 2× Euromedex PCR Master Mix (for PCR condition see **Supplementary Figure 1**). *COI* fragments were amplified using the primers COI_F106 and COI_R154 (**Supplementary Table 1**). Each primer contains a unique barcode sequence at the 5′ end. During amplification, the barcodes are integrated into the 3′ and 5′ ends of the amplicon, allowing 96 distinct barcode combinations (8F x 12R, **Supplementary Table 2**). The expected amplicon size was 750 bp. PCR products were pooled, purified, and prepared for sequencing using the SQK-LSK-114 kit (Oxford Nanopore Technologies, ONT). We performed sequencing on a MinION MK1C using Flongle flow cells (FLG-114).

Basecalling was conducted using the high-accuracy model in Dorado, and demultiplexing was performed with LAST (Frith et al., 2010; Hamada et al., 2017). Consensus sequences were generated using Medaka. Resulting *COI* sequences were queried against the NCBI nucleotide database (nt) using BLAST+ (blastn; Camacho et al., 2009) with the following parameters: - evalue 1e-5, -max_target_seqs 50, and -entrez_query "txid43845[Organism:exp]" to restrict searches to *Drosophilidae*. Searches were executed remotely on NCBI servers, and outputs were formatted in CSV. We imported BLAST results into R (version 2024.04.2+764) and filtered hits to retain only those with query coverage above 90%. Among these hits, we selected the highest percentage identity; in cases of ties, we retained the hit with the highest query coverage. When multiple equally scoring hits remained, we kept only one hit per taxon. Finally, because *COI* did not allow discrimination between *D. bipectinata* and *D. malerkotliana*, nor between *D. prosaltans* and *D. austrosaltans*, we identified these taxa as the *bipectinata* complex and the *saltans* subgroup, respectively.

### Microbiome analysis by metabarcoding

To assess microbiome composition across *Drosophila* species and localities, individuals were pooled in groups of ten, each comprising specimens from the same species and locality. When fewer than ten individuals were available, all available individuals were pooled, with the smallest pool consisting of three individuals. When insufficient numbers were available at the species level within a locality, individuals were pooled at the species-group level. Accordingly, “willistoni sp.” and “saltans sp.” refer to pools composed of individuals from multiple species within the *willistoni* and *saltans* species groups, respectively.

We performed DNA extraction using the ZymoBIOMICS DNA Miniprep kit following the manufacturer’s instructions. We included two controls: a negative extraction control, in which the sample was replaced with 750 µL of DNA/RNA Shield solution, and a positive control consisting of an artificial microbial community of known composition (ZymoBIOMICS D6300).

We amplified the bacterial 16S rRNA gene and the fungal ITS region, using distinct primer sets (**Supplementary Table 1;** see **Supplementary Figure 1** for PCR condition). Each primer was tagged with a unique synthetic barcode at the 5′ end. During amplification, the barcodes are integrated into the 3′ and 5′ ends of the amplicon, allowing sample identification after multiplexing (**Supplementary Tables 3 and 4**). In addition to extraction controls, PCR amplifications include a negative control consisting of nuclease-free water.

After amplification, we pooled 16S and ITS amplicons at a 90:10 ratio based on DNA concentration. We chose this ratio because the expected bacterial diversity is much higher than that of fungi. We performed library preparation as described previously for *COI* amplicons. Sequencing was conducted on a MinION MK1C using FLO-MIN-114 flow cells. Basecalling was performed directly on the MK1C using the high-accuracy model implemented in Dorado.

### Quality cleaning, clustering and taxonomic affiliation

Sequencing reads obtained from 16S and ITS amplicons were first separated by size (16S: ∼1350 bp; ITS: ∼450 bp) prior to demultiplexing using LAST (Frith et al., 2010; Hamada et al., 2017). Reads in which more than 20% of bases had a quality score below Q10 were considered low quality and removed using the fastq_quality_filter function from the FASTX-Toolkit (v0.0.14). Pools with low sequencing depth were subsequently excluded using thresholds of 1,000 reads for 16S and 400 reads for ITS. All remaining reads were then oriented in the same direction using the --orient option in VSEARCH (v2.30.0; Rognes et al., 2016).

Sequence clustering was performed using MeShClust v3 with default parameters (Girgis, 2022), generating clusters of similar sequences, each corresponding to an operational taxonomic unit (OTU). Clusters containing less than ten reads were discarded. The centroid sequence of each remaining cluster was retained to generate a FASTA file containing one representative sequence per OTU, and a contingency table was constructed to quantify read counts per OTU and per pool.

Chimeric sequences were identified and removed using the remove_chimera function in FROGS (v5.1.0) (Edgar et al., 2011; Rognes et al., 2016; Escudié et al., 2018). Taxonomic assignment of centroid sequences was performed using BLCA 2023 (Camacho et al., 2009; Gao et al., 2017), with the UNITE reference database (SH general release dynamic 19.02.2025) for ITS sequences and the SILVA database (SSURef NR99, release 138.2) for 16S sequences. Taxonomic ranks were assigned using a hierarchical probability threshold approach, as recommended for Bayesian classifiers (Gao et al., 2017). When species-level identification was not supported, OTUs were assigned to the lowest confidently supported rank and labelled with the suffix “sp.”. Sequences that could not be reliably assigned were classified as “Unclassified”.

In parallel, 16S centroid sequences were aligned using MAFFT (v7.525; Katoh and Standley, 2013), and a phylogenetic tree was inferred using FastTree (v2.2.0; Price et al., 2010) under a maximum-likelihood framework with the GTR model.

### Contamination Control and Data Filtering

To identify and filter potential contamination, reads detected in the three control samples were first examined. For each OTU, background read counts observed in controls were used to correct sample counts by subtracting these values from all samples containing the corresponding OTU. Data were Organised and processed using the phyloseq package in R. OTUs represented by fewer than two reads across the entire dataset were removed. In addition, OTUs corresponding to non-target taxa (i.e., non-bacterial sequences in 16S data and non-fungal sequences in ITS data) were excluded from further analyses. We also removed samples with insufficient sequencing depth. Based on rarefaction curves (**Supplementary Figures 2 and 3**), we selected read count thresholds to maximize sample retention while ensuring that most samples approached asymptotic richness. We set the minimum sequencing depth at 1,000 reads per sample for bacterial communities and 400 reads per sample for fungal communities.

### Statistical analyses

All statistical analyses were conducted in R (version 2024.04.2+764). Following *COI*-based identification, we calculated the number of specimens per host species for each locality, along with the proportion of invasive versus native individuals. Differences in IAS proportions among localities were tested using pairwise Fisher’s exact tests.

After 16S and ITS sequencing and data processing, alpha diversity was assessed for each *Drosophila* pool using species richness, the Shannon diversity index (H′), and Pielou’s evenness (J). We evaluated differences in alpha diversity across host species, host category (native vs. invasive), and locality using Kruskal–Wallis tests. When significant, we performed pairwise comparisons using Wilcoxon tests with Holm–Bonferroni correction.

We conducted beta diversity analyses separately for bacterial and fungal communities. For bacterial communities (16S), we assessed phylogenetic structure and composition using unweighted and weighted UniFrac distances (Lozupone and Knight, 2005). For fungal communities (ITS), Jaccard and Bray–Curtis distances were used to assess community composition and structure, respectively, because ITS sequences have high variability and alignment challenges.

We tested the effects of host species, locality, and host category using PERMANOVA. Homogeneity of multivariate dispersion was assessed using a multivariate analogue of Levene’s test (betadisper). When significant effects were detected, we conducted post hoc pairwise PERMANOVA comparisons with Bonferroni correction.

### Identification of candidate OTUs

OTUs assigned to the same species were collapsed by summing their read counts when reliable species-level identification was available. However, OTUs sharing the same lowest taxonomic assignment (e.g., “Genus sp.”) may represent distinct biological species. To avoid overinterpretation and artificial merging of potentially different taxa, we retained these OTUs as separate entities and assigned unique identifiers. Similarly, we treated unidentified OTUs (“Unclassified”) as distinct entities.

We then identified taxa exclusively associated with IAS or native species. Taxa identified above the species level may correspond to multiple species, including those identified elsewhere in the dataset. However, because all samples were processed using the same molecular marker and taxonomic pipeline, consistent failure to assign species-level identities was assumed to reflect true limitations in taxonomic resolution rather than annotation bias. Consequently, taxa identified at different taxonomic levels were treated as distinct entities, and taxonomic relationships were not used to infer identity when defining exclusivity.

For taxa identified only at the lowest confidently supported taxonomic level (e.g., “Genus sp.”), a conservative approach was adopted: all taxa sharing the same taxonomic label were considered collectively when assessing exclusivity. Such taxa were considered exclusive only if all corresponding OTUs were present in one category and absent from the other. We applied the same approach to unidentified taxa.

We defined the core bacteriome and mycobiome of each host species using a prevalence threshold of 50% and a relative abundance threshold of 0.001.

To place our results in a broader context, we applied the same analytical pipeline to three previously published datasets: Staubach et al. (2013), Wang et al. (2020), and Brown et al. (2023). For each dataset, we retained only wild-caught samples to ensure comparability. Specifically, we included 14 pools from North America (Staubach et al., 2013), 78 pools from the USA and Europe (Wang et al., 2020), and 190 individual flies from tropical Australia (Brown et al., 2023).

We reanalysed each dataset independently using the same pipeline as for our own data, then combined the results to generate a reference dataset representing the *Drosophila* microbiome across multiple geographic regions. We then quantified the abundance and prevalence of taxa identified as part of the core microbiome across host species and regions.

Because OTUs were inferred independently in each dataset, OTU identifiers could not be directly matched. Therefore, OTUs were collapsed at the species level where possible. For uncertain assignments (e.g., “Genus sp.”), taxa were collapsed at the lowest shared taxonomic level before comparing abundance and prevalence across datasets.

### *Drosophila* host tree reconstruction

Genomes assemblies (GCA_037075145.1,GCA_018904605.1,GCA_003401975.1, GCA_018150985.1, GCA_017639315.2, GCA_030179905.1, GCA_018153235.1, GCA_018150375.1, GCA_018903615.1, GCA_035045865.1, GCA_018151275.1, GCA_024703675.1, GCA_018151085.1, GCA_018903445.1, GCA_018150345.1, GCA_018151315.1) were downloaded from NCBI and analysed with BUSCO (v5.0.0, Seppey et al 2019). We identified 2740 genes as single orthologs present in all species and aligned them with MAFFT v7.505 (Katoh and Standley 2013). We cleaned alignments using TrimAl (Capella-Guttiérez et al. 2009). We randomly selected 203 alignments from the 2740 alignments and concatenated them into a supermatrix representing 139885 residues (33853 parsimony-informative residues). We reconstructed the tree using IQ-TREE3 (Wong et al. 2025, https://iqtree.github.io/) with the Q.INSECT+F+I+G4 model and UltrafastBootstrap. The resulting tree topology is congruent with the phylogeny in Suvorov et al. (2022), and all nodes are supported at 100%. We calibrated the tree using ultrametrization (R package ape; Paradis and Schliep 2018), with some nodes (identified by black circles) constrained within time ranges derived from data available in TimeTree5 (Kumar et al. 2022) and Suvorov et al. (2022). Ranges (minimum - maximum, in MYR) were (66-74) for *Scaptodrosophila (S.) latifasciaformis/D. melanogaster* most recent common ancestor (MRCA), and (56-62) for *Chymomyza costata/Zaprionus (Z.) indianus* MRCA.

## Results

### *Drosophila* community composition in French Guiana

We collected 744 *Drosophila* individuals, of which 719 were identified to species level using *COI* barcoding (**Figure 1A**). The remaining 25 individuals could not be reliably assigned due to the limited polymorphism of *COI*.

Eight native species were detected, belonging to the Neotropical *willistoni* and *saltans* species groups (**Figure 1B**). The *willistoni* species group comprised five species and was largely dominated by *D. paulistorum spp.* (95.67% of individuals), followed by *D. willistoni* (3.12%). The *saltans* species group included three taxa, with *D. sturtevanti* being the most prevalent (87.3%).

Five IAS were identified, including three from the melanogaster group: the bipectinata complex (*D. bipectinata* and *D. malerkotliana,* which could not be distinguished based on *COI* sequences), *D. ananassae*, and *D. melanogaster*. The bipectinata complex was the dominant IAS, accounting for 66% of invasive individuals.

The fourth IAS was *Zaprionus indianus*, which is phylogenetically distant from species of the melanogaster group. *Z. indianus* belongs to the tribe Drosophilini, and traditional classifications have recognized *Zaprionus* as a separate genus. However, recent phylogenetic studies indicate that this lineage is nested within the broader Drosophila clade, outside the subgenus Sophophora (Yassin et al., 2010; Commar et al., 2012). Although its phylogenetic position would have been informative for comparisons among host categories, insufficient replication after quality filtering precluded its inclusion in downstream analyses (only one 16S pool and no ITS pools remained).

Finally, we identified specimens of *Scaptodrosophila latifasciaeformis*, a phylogenetically distant species, which were excluded from downstream analyses resulting in a final set of three IAS.

IAS prevalence strongly varied across localities (**Figure 1A**). Cayenne was largely dominated by IAS (91.2%), whereas proportions were significantly lower in Kaw (6.4%), Bélizon (0.4%), and the Nouragues (2.2%) (all p < 0.001 vs. Cayenne). Among remote sites, IAS frequencies differed only between Kaw and Bélizon (p = 1.37 × 10⁻³).

IAS diversity followed a similar pattern: all IAS species were recorded in Cayenne, compared to two in Kaw and only one in Bélizon and the Nouragues (**Figure 1A**). The *bipectinata complex* was present in all localities, whereas only *D. melanogaster* was detected outside Cayenne (in Kaw).

Although we occasionally detected IAS in forested sites, they remained largely confined to anthropised environments. IAS were therefore largely restricted to anthropised environments, prompting us to investigate whether microbiome composition differed between IAS and native species along this gradient.

### Quantification of diversity of microbial communities associated with *Drosophila* species in French Guiana

We first quantified bacterial and fungal diversity across locality, host species and host categories. To do so, we analysed 46 pools of native species and 16 pools of IAS, each comprising about 10 individuals from the same species and locality (**Supplementary Figure 4**). We extracted DNA from each pool, and amplified the bacterial 16S rRNA gene and the fungal ITS region before sequencing using Nanopore technology.

Rarefaction curves indicated sufficient sequencing depth to capture the most abundant taxa (**Supplementary Figures 2 and 3**).

We detected a total of 332 OTUs in the bacteriome, including 206 shared between native species and IAS (**Figure 2A**). Only 21 OTUs were unique to IAS, compared to 105 in native species. For the mycobiome, we identified 70 OTUs, of which 57 were shared; 13 were unique to native species, and none were specific to IAS (**Figure 2B**).

**Figure 2.**
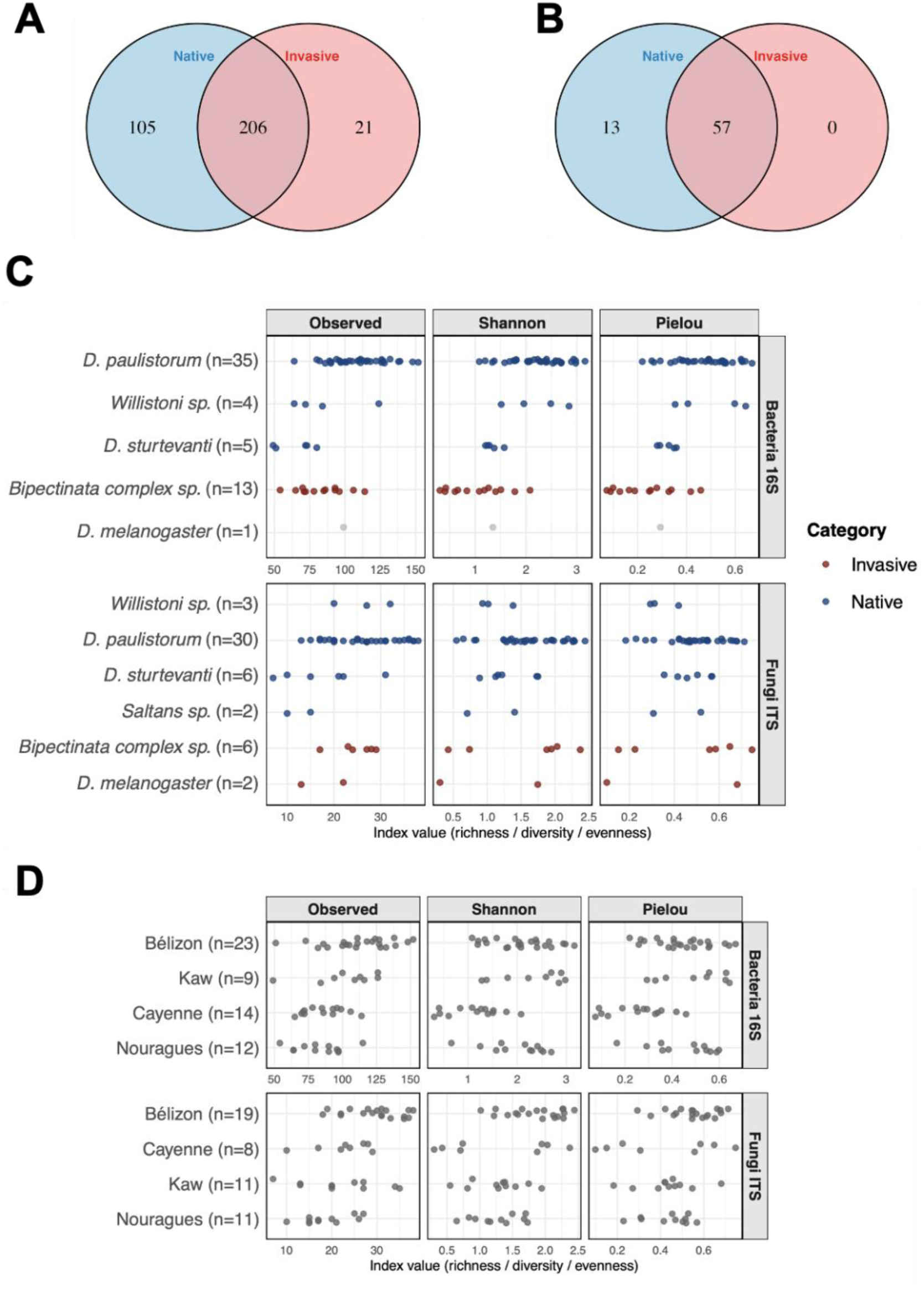
Alpha diversity of microbial communities associated with *Drosophila* species in French Guiana. (A–B) Venn diagrams showing the overlap of OTUs between invasive and native species for bacteria (A) and fungi (B). (C) Diversity indices by host species and (D) by sampling locality. Observed richness (left), Shannon diversity (centre), and Pielou’s evenness (right) are shown. Colors indicate host status, with IAS in red and native species in blue. For each panel, the upper row corresponds to bacterial communities (16S) and the lower row to fungal communities (ITS). In panel C, grey points indicate species with insufficient sample size for statistical analysis. Sample sizes (n) correspond to the number of pools, each consisting of 3 to 10 flies of the same species collected at the same site.

Bacterial alpha diversity (richness, Shannon index, and Pielou’s evenness) differed significantly among host species, locality, and “invasiveness” host category (**Table 1; Figure 2C**). However, pairwise comparisons revealed no clear pattern for richness across groups. In contrast, Shannon diversity and Pielou’s evenness were consistently lower in Cayenne and in the *bipectinata* complex, indicating microbiomes composed of fewer dominant taxa and a less even distribution of bacterial abundances compared to other localities and host species (**Table 1; Figure 2C; Supplementary Table 5**).

**Table 1:** Significant Kruskal-Wallis for observed specific richness and Shannon index.

| Dataset | Model | Term | P-value |
| --- | --- | --- | --- |
| Bacteriome | Richness | HostSpecies | 4.29E-05 |
|  |  | Category | 0.010387 |
|  |  | Locality | 0.000543 |
|  | Shannon | HostSpecies | 6.78E-06 |
|  |  | Category | 7.82E-06 |
|  |  | Locality | 0.000346 |
|  | Pielou | HostSpecies | 1.63E-05 |
|  |  | Category | 8.51E-06 |
|  |  | Locality | 0.000455 |
| Mycobiome | Richness | Locality | 0.00112 |
|  | Shannon | Locality | 0.005896 |
|  | Pielou | Locality | 0.043054 |

For fungi, host species and category had no significant effect on alpha diversity, which was instead driven by locality (**Figure 2D; Table 1**). Diversity was lower in Nouragues and Kaw than in Bélizon, indicating variation among rainforest sites even within similar host assemblages (**Figure 2D; Table 1; Supplementary Table 5**).

In conclusion, bacteriome diversity differed significantly according to host species, locality, and host category, whereas the mycobiome is primarily structured by locality.

### Microbiome community structure analysis in *Drosophila* from French Guiana

We further explored the complexity of microbial communities and structure to better evaluate their potential impact on IAS and native species (**Figure 3; Table 2**).

**Figure 3.**
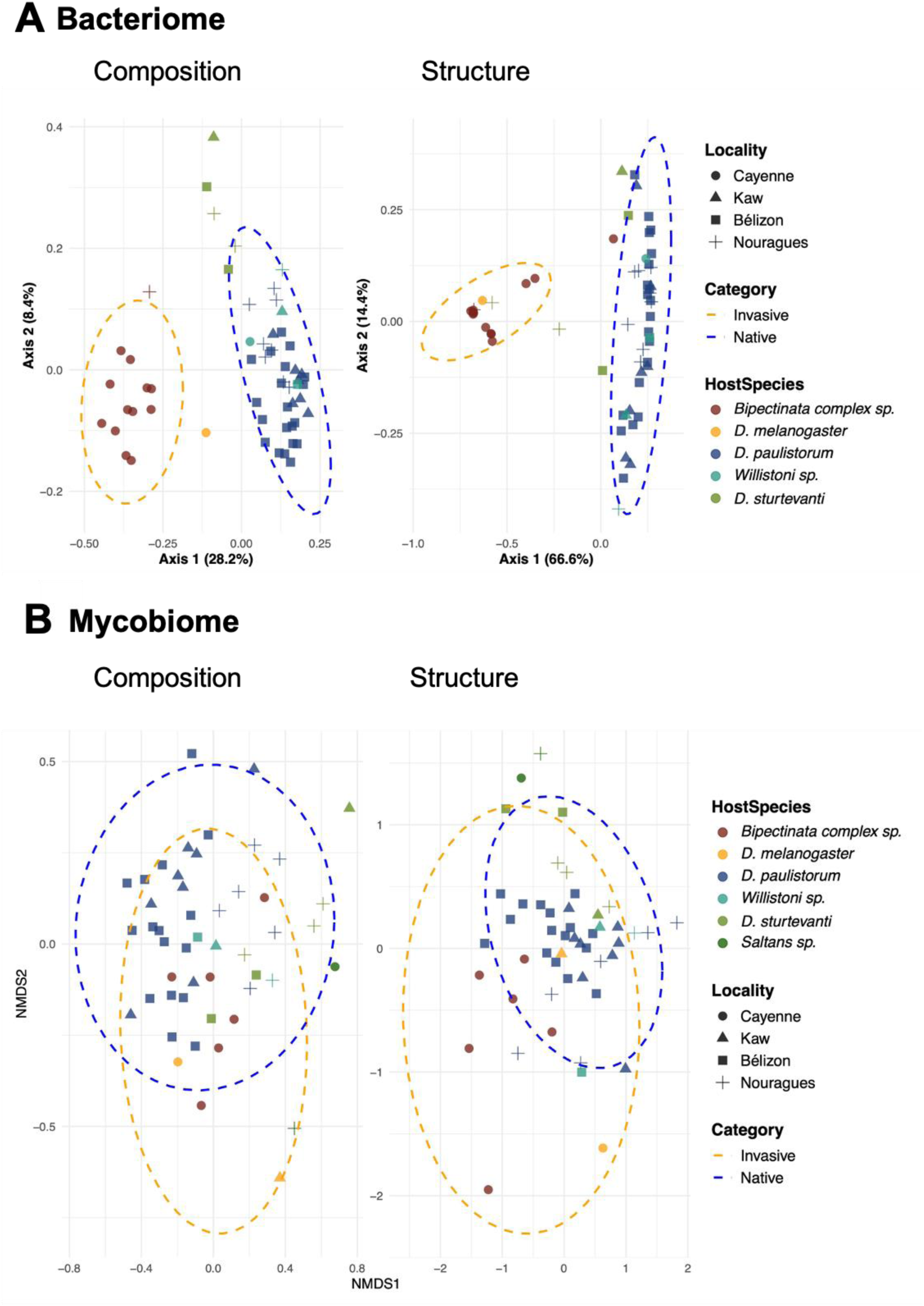
Ordination plots for bacteriome (A) and mycobiome (B). This graphic shows a PCoA based on composition (left) and structure (right) dissimilarities for the bacteriome (A); and a NMDS based on composition (left) and structure (right) dissimilarities for mycobiome (B). Each point represents a microbial community sampled from a pool of 10 flies. Point shape indicates the collection site (Locality), and point color indicates the host species (HostSpecies). The distribution of pools according to their category (IAS or native species) is indicated by dashed ellipses.

**Table 2.** Significant Permanova results for ordination analyses.

| Dataset | Model | Test | Term | P-value | R2 |
| --- | --- | --- | --- | --- | --- |
| Bacteria | UniFrac | ~ HostSpecies X Locality | HostSpecies | 0.001 | 0.16 |
|  |  |  | Locality | 0.001 | 0.27 |
|  |  |  | Locality:<br>HostSpecies | 0.031 | 0.08 |
|  |  | ~ Locality X Category | Category | 0.001 | 0.04 |
|  |  |  | Locality | 0.001 | 0.27 |
|  |  | ~ Category X Locality | Category | 0.001 | 0.25 |
|  |  |  | Locality | 0.005 | 0.07 |
|  | Weighted<br>UniFrac | ~ HostSpecies X Locality | HostSpecies | 0.001 | 0.19 |
|  |  |  | Locality | 0.001 | 0.45 |
|  |  |  | Locality:<br>HostSpecies | 0.026 | 0.06 |
|  |  | ~ Locality X Category | Category | 0.001 | 0.11 |
|  |  |  | Locality | 0.001 | 0.45 |
|  |  | ~ Category X Locality | Category | 0.001 | 0.53 |
| Fungi | Bray-Curtis | ~ HostSpecies X Locality | HostSpecies | 0.001 | 0.16 |
|  |  |  | Locality | 0.001 | 0.17 |
|  |  | ~ Locality X Category | Locality | 0.001 | 0.17 |
|  | Jaccard | ~ HostSpecies X Locality | HostSpecies | 0.001 | 0.13 |
|  |  |  | Locality | 0.001 | 0.13 |
|  |  |  | Locality:<br>HostSpecies | 0.043 | 0.1 |
|  |  | ~ Locality X Category | Category | 0.142 | 0.02 |
|  |  |  | Locality | 0.001 | 0.13 |
|  |  |  | Locality:<br>Category | 0.001 | 0.04 |

For the bacteriome, community composition and structure differed significantly according to host category, host species, and locality (**Figure 3A; Table 2**). Host category explained the largest proportion of variation among the factors examined; however, it was strongly confounded with host species and locality. Pairwise comparisons indicated greater similarity among species within the same species group. In contrast, stronger differences were observed between species groups, a pattern consistent with the influence of host phylogenetic relatedness. Microbiomes were similar among species within the same species group (e.g., willistoni or saltans). In contrast, clear differences were observed between species belonging to different groups, particularly between the willistoni and saltans groups and the bipectinata complex (**Figure 3A, Supplementary Table 6**). Locality further seems to influence bacteriome variation (**Supplementary Table 6**).

In contrast, mycobiome composition and structure were not affected by host category but varied significantly with host species and locality (**Figure 3B; Table 2**). Differences were primarily observed between species groups (*willistoni* species group vs. *saltans* species group), whereas species within the same group showed limited differentiation.

Most localities differed significantly in mycobiome composition and structure, except Kaw (**Supplementary Table 6**).

Overall, the mycobiome showed no significant differences according to host category, whereas significant variation was associated with locality and host species. Differences among host species were mainly observed between species groups, consistent with a possible contribution of host evolutionary relationships. However, the absence of a significant effect of host category on fungal communities does not preclude finer-scale differences. To better characterise the taxonomic basis of these patterns, we examined microbiome composition at the class level.

### Microbiome taxonomic overview at class level

OTUs were assigned to 310 bacterial taxa, including 296 detected in native species and 210 in IAS, of which 196 were shared between both categories. Similarly, 50 fungal taxa were identified, all of which were detected in native species, while 42 were additionally detected in IAS.

The bacteriome comprised *Alphaproteobacteria*, *Gammaproteobacteria*, *Bacilli*, *Bacteroidia*, and *Clostridia* (**Figure 4A**). *Alphaproteobacteria* were generally more abundant in IAS pools, whereas *Gammaproteobacteria*, *Bacilli*, and *Bacteroidia* were more represented in native hosts. However, three pools deviated from this pattern, including one *bipectinata complex* pool from Cayenne dominated by Gammaproteobacteria and two *D. sturtevanti* pools from the Nouragues showing elevated *Alphaproteobacteria* abundance (asterisk in **Figure 4A**). Bacterial class-level composition did not show any consistent association with host species or locality.

**Figure 4.**
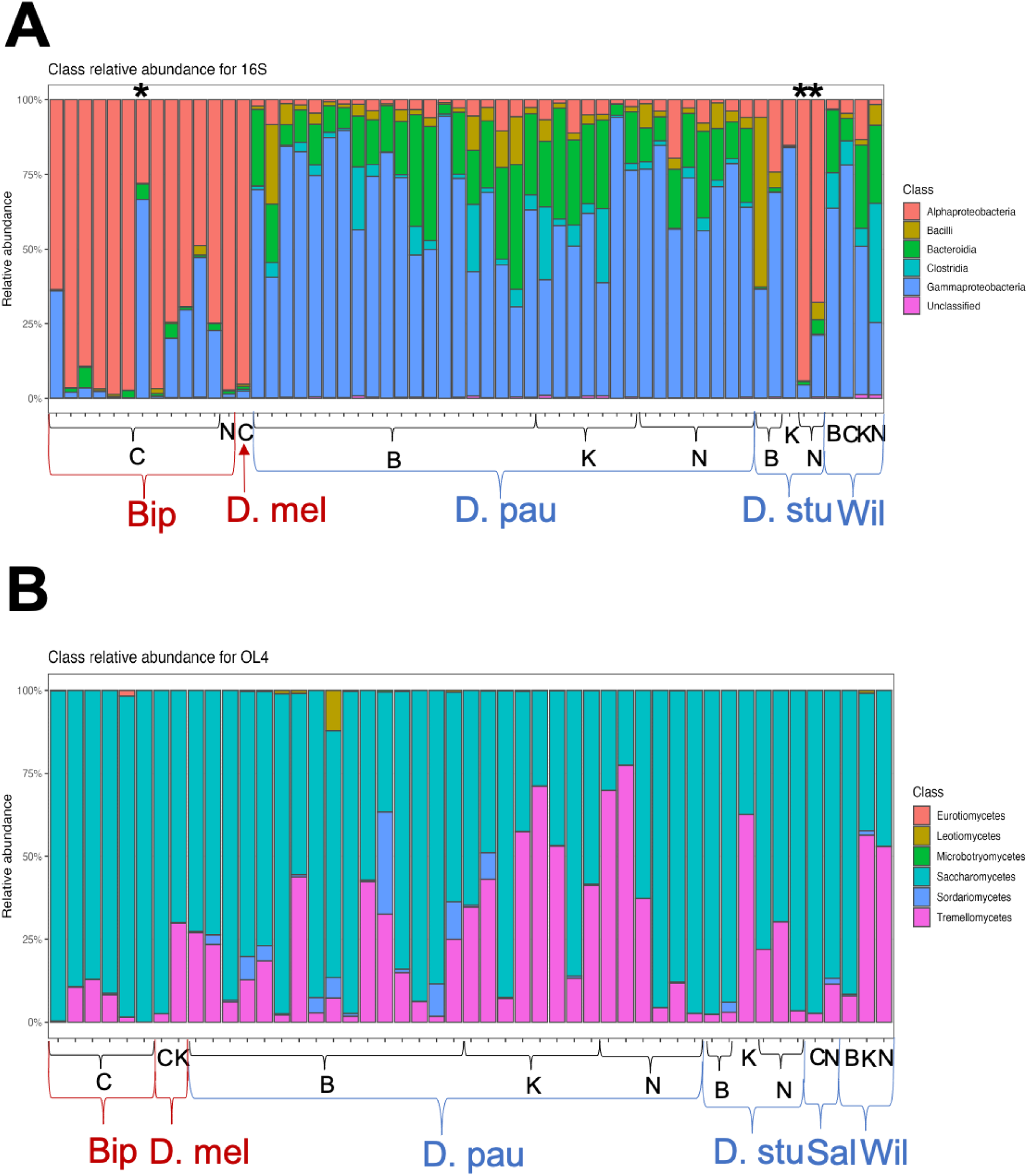
Barplot showing the relative abundance of bacterial classes across pooled fly bacteriomes (A) and mycobiome (B). Each bar represents a pool of 10 individuals. Host species are indicated on the x-axis (Bip: *Bipectinata* complex; D. mel: *D. melanogaster*; D. pau: *D. paulistorum*; D. stu: *D. sturtevanti*; Wil: *willistoni* species group; Sal: *saltans species group*). IAS are shown in red and native species in blue. Sampling localities are indicated by black labels (C: Cayenne; N: Nouragues; B: Bélizon; K: Kaw). Colors within bars represent the relative abundance of each bacterial (A) and fungi class (B). The asterisks indicate the pools whose class composition differs.

Fungal communities were largely dominated by *Saccharomycetes* across both IAS and native species, with *Tremellomycetes* (*basidiomycetes*) as the second most abundant class. Other fungal classes, *Eurotiomycetes*, *Leotiomycetes*, *Microbotryomycetes*, and *Sordariomycetes* (all *ascomycetes*) were sporadically detected in some pools. In contrast to bacteria, no clear pattern differentiates IAS from native taxa at the class level (**Figure 4B**).

To identify the taxa underlying these patterns, we next examined microbiome composition at a finer taxonomic resolution.

### Bacteriome taxonomic composition

We first identified bacterial taxa exclusively associated with either native hosts or IAS (**Supplementary Table 7**). Because some OTUs could not be assigned at the species level, the lowest available taxonomic rank was retained (“sp.”), with unique identifiers distinguishing OTUs sharing the same affiliation (see Materials and Methods for details).

We identified 2 taxa exclusive to IAS, both restricted to the *bipectinata* complex and characterised by low relative abundance but relatively high prevalence. In contrast, 15 taxa were exclusive to native hosts, displaying a wide range of prevalence and abundance (**Supplementary Table 7**). Only 1 unidentified *Candidatus Soleaferrea* among the 17 exclusive bacteria exhibited both high relative abundance and high prevalence (**Supplementary Table 7**).

These results further support the lower bacterial diversity observed in IAS and highlight clear compositional differences between native and invasive bacteriomes. These taxa best describe the microbiome composition differences between native species and IAS. These taxa therefore represent candidate bacterial taxa distinguishing the microbiomes of native and IAS hosts and were retained for further investigation.

To further investigate differences in host–microbiome associations, we defined the core bacteriome for host species represented by multiple pools using a minimum prevalence threshold of 0.5 and a relative abundance threshold of 0.001 (**Figure 5A**). The core bacteriome of the *bipectinata* complex comprised only 8 taxa, whereas native species harboured between 13 and 18 taxa, consistent with the lower bacterial diversity observed in IAS pools (**Figure 5A**).

**Figure 5.**
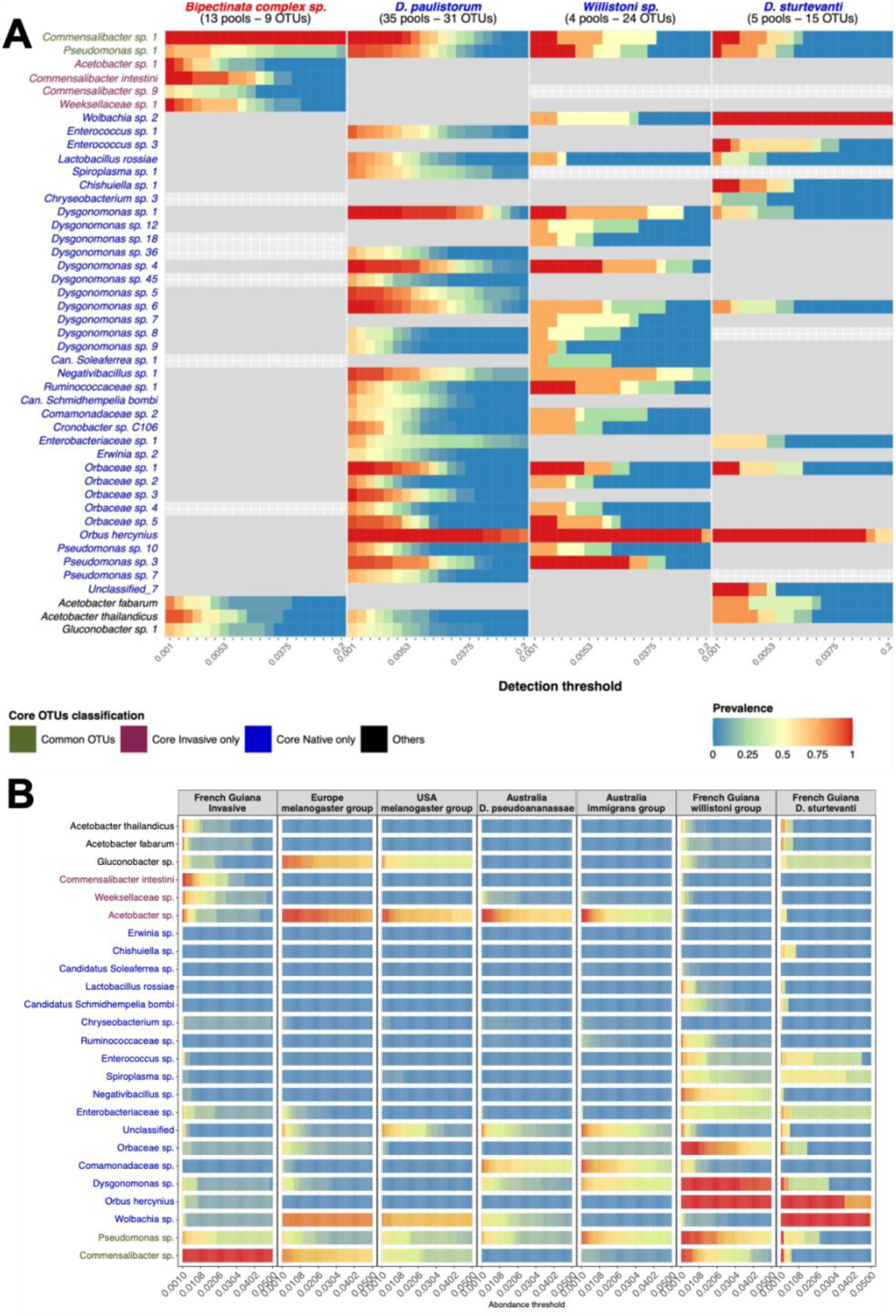
Heat map of the core bacteriome (A) and heat map of abundance and prevalence of core taxa in the *Drosophila* microbiome reference dataset (B). In A, each panel represents the core bacteriome of a host species. The x-axis indicates the abundance threshold, the y-axis lists the taxa names, and color represents prevalence across the number of pools indicated next to the host species name. Each colored square therefore indicates the prevalence (color scale) of a given taxon (y-axis) at a given abundance threshold (x-axis) for a given host species (panel). Grey cells indicate that the taxon was detected in the corresponding host but did not meet the core criteria. Taxa on the y-axis are colored according to their distribution among host species: red indicates taxa found in the core bacteriome of IAS but not in native species; blue indicates taxa found in the core bacteriome of native species but not in IAS; black indicates taxa found in some native and some IAS but not in all host species; green indicates taxa found in all host species. In B, facets refer to a single host at a single locality. The color legend used is the same as for A.

Two taxa were found across all host species, while 3 additional taxa were shared between IAS and some native species but were not ubiquitous, suggesting they are unlikely to drive differences between host categories (**Figure 5A**). In contrast, 19 taxa were present in the core bacteriome of native species but absent from IAS, whereas only 4 taxa were present in the IAS core bacteriome but absent from native core bacteriomes (**Figure 5A**). These marked asymmetries indicate strong differences in host–symbiont associations between native and IAS. Furthermore, *D. paulistorum* and *Willistoni* sp., both belonging to the *willistoni* species group, were more similar to each other than either was to *D. sturtevanti* (from the *saltans* species group), indicating that shared bacterial taxa differ among *Drosophila* species groups. This result suggests that bacteriome composition also reflects host phylogenetic relationships (**Figure 5A**).

Those 19 native-core and 4 IAS-core taxa that distinguished the core bacteriomes of the two host categories were therefore retained as candidates for future taxonomic and functional investigation. If they are functionally different, they could strongly impact invasion dynamics; therefore, we retained as candidate taxa potentially involved in invasion dynamics. However, given the lack of available information, Unclassified 7 was not retained as a candidate taxon. *Candidatus Soleaferrea* is already in the candidate list because we found it exclusively in native hosts.

Among both specific and core bacterial taxa, 52 OTUs could not be identified at the species level and were therefore reported at the genus or higher taxonomic rank. Among these, 5 taxa are unidentified at the species level because the 16S rRNA marker did not provide sufficient resolution to distinguish closely related species. The remaining unidentified taxa at species level showed no sufficiently similar matches in reference databases, suggesting that they may correspond to previously undescribed local bacterial species.

To place these results in a broader context, we compared core taxa with a global reference dataset representing the native *Drosophila* bacteriome across multiple geographic regions worldwide, including the USA, Europe, and tropical Australia (for more details, see Materials and Methods and Staubach et al., 2013; Wang et al., 2020; Brown et al., 2023; **Figure 5B**).

Some taxa, such as *Acetobacter* sp., which were detected as core members of the microbiome in IAS but not in native species, were consistently abundant across regions, indicating that this genus constitutes a widespread component of the *Drosophila* microbiome.

The absence of *Acetobacter* sp. from the core microbiome of native Guianese species therefore suggests that these native species groups may not have a particular affinity for this bacterium, despite its broad distribution across hosts in other regions worldwide.

In contrast, *Commensalibacter intestini* and an unidentified Weeksellaceae were restricted to neotropical French Guiana, where they reached high abundance and prevalence only in IAS, suggesting a strong local association with IAS *Drosophila* (**Figure 5B**).

However, this pattern appeared to be specific to *C. intestini*, as other *Commensalibacter* taxa were consistently abundant and prevalent across regions, supporting the idea that the genus *Commensalibacter* constitutes a widespread core component of the *Drosophila* microbiome (**Figure 5B**).

Moreover, 13 of the 17 taxa present in the native core microbiome but absent from the IAS core microbiome were not detected as core taxa in the reference dataset (**Figure 5B**), indicating that they are locally restricted and specifically associated with native hosts.

Among the 4 remaining taxa, unidentified *Comamonadaceae* and unidentified *Dysgonomonas* were abundant and prevalent only in tropical Queensland (Australia); the unclassified group, which gathers a lot of completely different OTUs was ubiquitous across regions, and *Wolbachia* showed a host-dependent pattern, as datasets from Europe and the USA are based solely on *D. melanogaster* (**Figure 5B**).

Overall, these results revealed a pronounced asymmetry between native and IAS hosts, with native species harbouring a richer and more diverse core microbiome. Based on exclusivity, abundance/prevalence, and global distribution patterns, we identified 38 bacterial taxa that distinguished native and IAS microbiomes and retained them as candidates for future taxonomic and functional investigation.

### Mycobiome taxonomic composition

Although no significant differences in mycobiome diversity, composition, or structure were detected between IAS and native species, we looked for taxa exclusively associated with each host category (**Supplementary Table 7**). We found 8 fungal taxa exclusive to native species, whereas none were exclusive to IAS. These taxa showed variable prevalence and abundance, suggesting heterogeneous but potentially ecologically relevant associations with native hosts; therefore, they were retained as candidate taxa distinguishing native and IAS hosts at finer taxonomic resolution.

Next, we defined the core mycobiome for host species represented by multiple pools (**Figure 6**). In contrast to the bacteriome, core mycobiome richness was similar between IAS (5–12 taxa) and native species (6–11 taxa), indicating limited differences in overall fungal diversity. 3 taxa were shared across all host species, while 4 additional taxa were shared between IAS and some native species but were not ubiquitous (**Figure 6**). These 7 taxa are therefore unlikely to contribute to differences between host categories.

**Figure 6.**
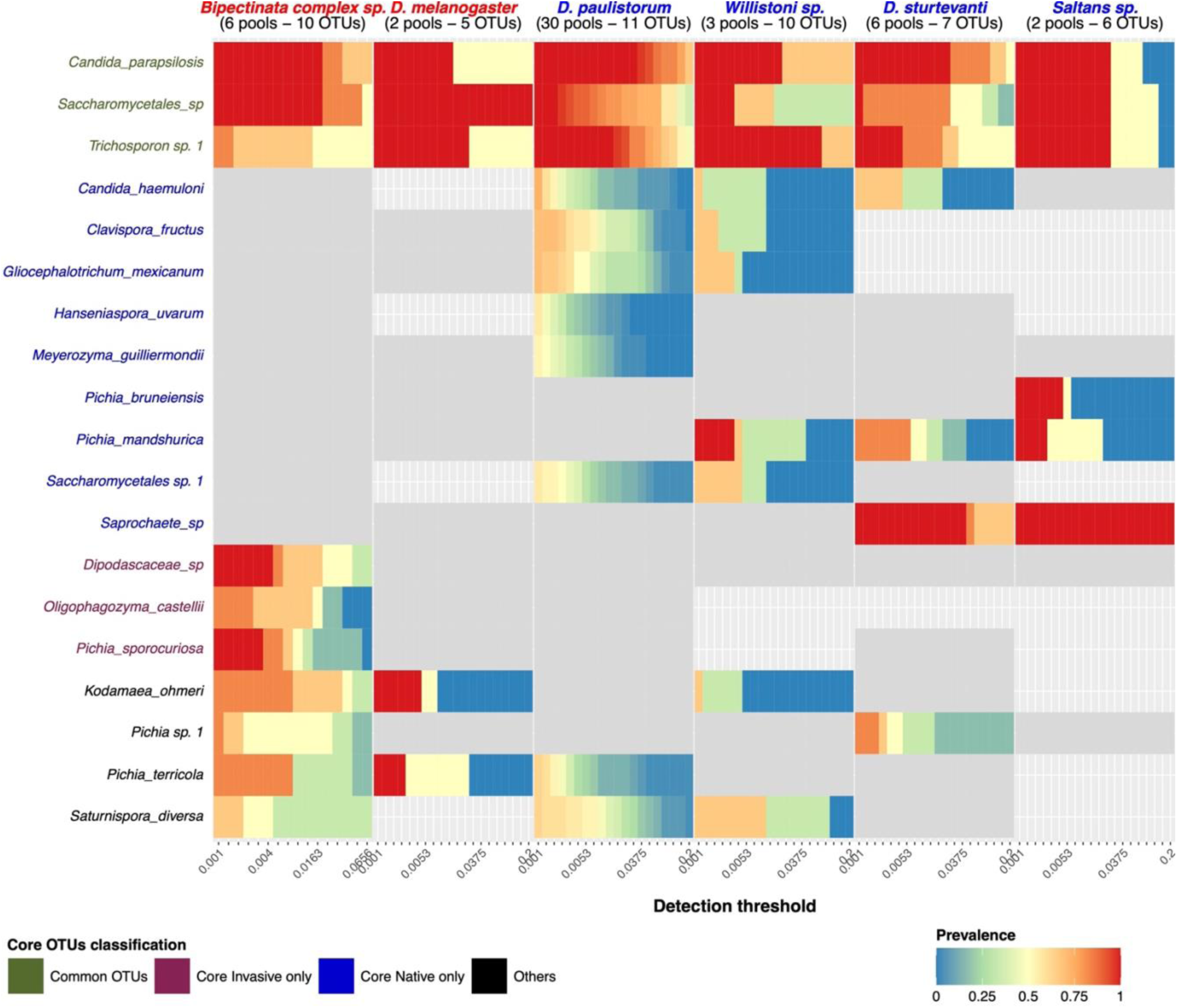
Heat map of the core mycobiome. Each panel represents the core mycobiome of a host species. The x-axis indicates the abundance threshold, the y-axis lists the taxa names, and color represents prevalence. Each colored square therefore indicates the prevalence (color scale) of a given taxon (y-axis) at a given abundance threshold (x-axis) for a given host species (panel). Grey cells indicate that the taxon was detected in the corresponding host but did not meet the core criteria. Taxa on the y-axis are colored according to their distribution among host species: red indicates taxa found in the core mycobiome of IAS but not in native species core mycobiome; blue indicates taxa found in the core mycobiome of native species but not in IAS core mycobiome; black indicates taxa found in some native and some IAS but not in all host species; green indicates taxa found in all host species.

Nevertheless, 9 taxa were present in the core mycobiome of native species but absent from the IAS core, whereas only 3 taxa were specific to the IAS core mycobiome (**Figure 6, Table 3**). We retained these 12 taxa as candidates potentially involved in invasion dynamics. While the number of IAS-specific core fungal taxa was comparable to that observed in bacteria, native-specific core taxa were markedly less numerous in fungi than in bacteria.

**Table 3.**
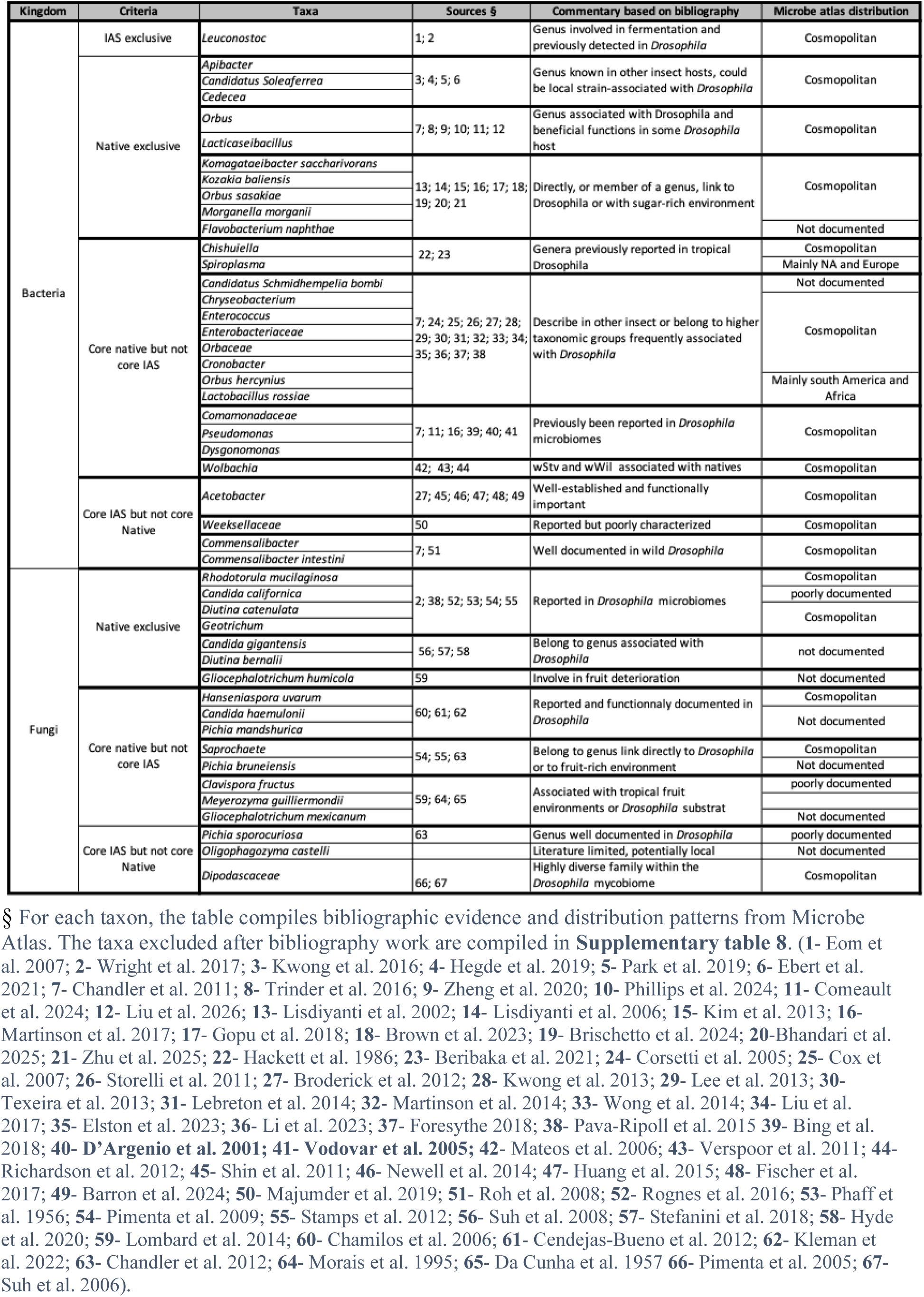
Summary of candidate taxa based on criteria, literature review and global distribution. Taxa are reported at the lowest available taxonomic level. OTUs sharing the same taxonomic assignment and the same point of interest are grouped.

Overall, differences between IAS and native core communities therefore appeared less pronounced in the mycobiome than in the bacteriome, although some taxa differed in their occurrence among host categories.

Patterns of taxon sharing between host species groups were relatively similar, with only moderate differences between the willistoni and saltans groups, suggesting a weaker influence of host phylogeny on fungal communities compared to bacterial communities.

Among both specific and core fungal taxa, 8 OTUs could not be identified at the species level because no sufficiently similar matches were found in reference databases, a pattern consistent with weaker phylogenetic structuring of fungal than bacterial communities.

Overall, in contrast to the bacteriome, the fungal core microbiome appears broadly similar across host categories and phylogenetic groups. However, 8 taxa were exclusively detected in native hosts, 9 others were highly abundant and prevalent only in native species, and 3 taxa were identified as core exclusively in IAS. These 20 fungal taxa represent candidates distinguishing the mycobiomes of native and IAS hosts and provide targets for future taxonomic and functional investigation.

## Discussion

### *Drosophila* community in French Guiana and invasion status

All eight native Neotropical Drosophila species we have sampled in French Guiana belong to two closely related lineages, the willistoni and saltans species groups, which diverged from the melanogaster group more than 40 million years ago and are well documented across Latin America (Burla et al., 1949; Tamura et al., 2003; O’Grady and DeSalle, 2018), whereas the three IAS (the *bipectinata* complex, *D. ananassae* and *D. melanogaster*) belong to the melanogaster species groups. In addition to the IAS previously mentioned, individuals identified as *Z. indianus* and *Sc. latifasciaeformis* were also captured; however, neither species was included in the subsequent analyses.

The *bipectinata* complex refers to *D. bipectinata* and *D. malerkotliana*, which cannot be clearly distinguished based on *COI* sequences (Kopp and Barmina, 2005). However, since only *D. malerkotliana* is recognized as IAS in Latin America, particularly in Brazil, it is likely the species corresponding to the specimens detected here (Yuzuki and Tidon, 2020). Both the *bipectinata* complex and *D. ananassae* belong to the *ananassae* subgroup and are cosmopolitan species native to Southeast Asia (Das et al., 2004; Matsuda et al., 2009). *D. melanogaster*, originally from Africa, has spread globally via Europe and reached the Americas during the transatlantic slave trade (Kao et al., 2015; Dutheil, 2020). Overall, the presence of the sampled species in French Guiana is consistent with their known distributions in different parts of South America like Brazil and Ecuador (Acurio et al. 2010; Yuzuki and Tidon, 2020).

The relatively low capture rate (744 individuals from 556 traps) may be explained by the cotton barrier and rain cover used to protect microbiome integrity, which likely reduced bait attractiveness by limiting volatile diffusion. Nevertheless, rarefaction curves and comparison with previous records indicate that a substantial proportion of the *Drosophila* diversity detectable with our sampling method was captured (Miller and Hua-Van, unpublished data since 2014; Yuzuki and Tidon, 2020; Baião et al., 2023; Prediger et al., 2024; Papachristos et al., 2025; **Supplementary Figure 5**).

In our study, we documented an increase in the proportion and diversity of IAS along the anthropisation gradient, consistent with previous studies (Francis and Chadwick, 2015; Potgieter et al., 2024; Ulmer et al., 2024; Kapun et al., 2026). IAS remain extremely rare in rainforest localities such as Nouragues and Bélizon, despite their expected range expansion (Sakai et al., 2001; Arim et al., 2006). Their occasional presence in forested sites indicates that dispersal to these localities occurs, whereas their low frequency suggests that environmental or ecological conditions may limit their persistence. Such limitations in IAS expansion may rely not only on genetic traits and their behaviour but also on the microbiome as an extended phenotype (Calderon-Cortés et al., 2012; Davenport et al., 2017; Kolodny et al., 2020; Henry et al., 2021; Petersen et al., 2023; Bordenstein et al., 2024).

Overall, while IAS are established in French Guiana, their distribution remains largely concentrated in coastal anthropised habitats, whereas native species dominated rainforest localities. This contrasting distribution provides an ecological context in which to compare the microbiomes associated with native and invasive Drosophila.

### Diversity, composition and structure differences between invasive and native species

In this study, bacteriome diversity, composition, and structure differed according to host species, host category, and locality. In particular, bacterial diversity was lower in IAS, while bacterial communities differed between IAS and native hosts, alongside variation associated with host species and locality.

Host species and geographic location are known to be major determinants of Drosophila bacteriome composition and structure (Chandler et al., 2011; Campbell et al., 2012; Staubach et al., 2013; Wang et al., 2020; Wang et al., 2022; Ying et al., 2022; Brown et al., 2023). It is well established that Drosophila acquire most of their bacteriome from the environment (Chandler et al. 2011; Broderick and Lemaitre 2012; Staubach et al. 2013; Wang et al. 2020). But, hosts themselves likely act as selective filters determining which microorganisms can persist (Broderick and Lemaitre 2012; Douglas 2018; Ma and Leulier, 2018; Pais et al., 2018). In this context, Baas Becking’s conjecture, “everything is everywhere, but the environment selects” is particularly relevant (Baas Becking, 1934).

Previous studies suggest that IAS microbiomes are shaped by both microbial loss during introduction events (Torchin et al., 2003; MacLeod et al., 2010; Szklarzewicz et al., 2020; Mowery et al., 2024; Romeo et al., 2025) and subsequent acquisition of local microorganisms (Christian et al., 2018; Santos et al., 2021; Chiarello et al., 2022; Zhu et al., 2022; Do et al., 2023; Grbin et al., 2023; Maritan et al., 2024; Zuo et al., 2024). Rare microbial taxa may be lost during introduction and therefore no longer vertically transmitted in subsequent generations, whereas local microorganisms may be horizontally acquired depending on host affinity. As a result, the composition and structure of IAS microbiomes are expected to differ from those of native species, because vertically inherited bacteria partly reflect the native range of IAS (Mateos et al. 2006; Richardson et al. 2012; Szklarzewicz et al., 2020), while newly acquired taxa are filtered by the local environment and host species or phylogeny (Broderick and Lemaitre 2012; Douglas 2018; Ma and Leulier, 2018; Pais et al., 2018). In contrast, the direction of diversity change depends on the balance between taxa lost and taxa acquired: microbiome diversity may decrease when losses exceed acquisitions (Goddard-Dwyer et al., 2021; Santos et al., 2021; Zhu et al., 2022; Li et al., 2022; Zuo et al., 2024; Romeo et al., 2025). In contrast, it may increase when acquisition of local microorganisms exceeds taxon loss (Stevens and Olson, 2013; Duguma et al., 2017; Chiarello et al., 2022; Romeo et al., 2025).

Our comparison with reference datasets revealed geographic variation consistent with this general framework. Some taxa, such as *Acetobacter thailandicus* and *Commensalibacter intestini*, were detected in our dataset only in IAS from French Guiana. This restricted distribution is compatible with local acquisition, although the present data cannot distinguish acquisition in the invaded range from other explanations, including geographic variation in microbiome composition or incomplete detection in reference datasets. In contrast, taxa such as *Pseudomonas* and *Acetobacter* were detected across multiple geographic regions and host species, consistent with their widespread association with Drosophila (**Figure 5B**).

Anthropised environments are also known to influence microbiome diversity and structure (Yan et al., 2016; Parajuli et al., 2018; Dillard et al., 2022; Maraci et al., 2022; Li et al., 2023; Magura et al., 2024; Nguyen et al., 2024; Yan et al., 2024). Because IAS were detected almost exclusively in Cayenne, the only anthropised locality in our study, the respective effects of locality and host category cannot be clearly disentangled. Moreover, although bacteriome differences between individual host species were relatively weak, stronger differences were observed between more distantly related species groups (e.g., willistoni vs. saltans), suggesting an influence of host phylogeny. Consequently, the respective contributions of host category, environmental conditions, and host evolutionary history associated with the observed bacteriome differences cannot be fully separated in the present sampling design.

In contrast, for the mycobiome, we found that community composition and structure differed only among host species and localities, whereas diversity was influenced solely by locality. This pattern is consistent with previous studies showing that fungal communities are more environmentally driven than bacterial ones, with host species generally acting as a secondary factor (Huffnagle and Noverr, 2013; Sun et al., 2021; Worsley et al., 2022; Dragičević et al., 2023; Medeiros et al., 2025). In our study, the host species effect appeared to be phylogenetically structured, as differences were detected only between host species belonging to distinct species groups, suggesting a relatively limited influence of the host species. The absence of significant differences in microbiome composition and structure between IAS and native host does not exclude finer taxon-specific differences, which were subsequently examined through core and exclusive taxa. Whether such differences have functional consequences for the host remains to be experimentally determined.

Finally, statistical evidence indicates that bacteriome diversity is reduced in IAS and that its composition and structure differ markedly from those of native species. Overall, bacterial communities differed between IAS and native hosts, whereas fungal communities showed stronger associations with locality and host species. Because 16S and ITS metabarcoding provide primarily taxonomic rather than functional information, the consequences of these compositional differences for host phenotype cannot be determined from the present data.

Finer taxonomic and functional approaches are therefore required to characterise the biological significance of the taxa identified here.

### General microbiome taxonomic composition

*Alphaproteobacteria* and *Gammaproteobacteria* were the two dominant classes in the bacteriome, in IAS and native species, respectively. *Bacilli*, *Bacteroidia*, and *Clostridia* were primarily found in native species and were less frequent in IAS, indicating that differences in bacteriome composition and structure between IAS and native species are detectable even at high taxonomic levels. These patterns are consistent with previous studies reporting *Alphaproteobacteria* as the dominant class in *Drosophila*, with *Gammaproteobacteria* also prevalent depending on environmental conditions, while other classes are typically present at lower abundance (Chandler et al., 2011; Staubach et al., 2013; McMullen et al., 2021; Brown et al., 2023; Cho and Kang, 2025).

Two pools of *D. sturtevanti* from the Nouragues and one *bipectinata* complex pool from Cayenne deviated from the general class-level composition patterns of native and IAS, showing unusually high abundances of *Alphaproteobacteria* and *Gammaproteobacteria*, respectively. In *D. sturtevanti*, this pattern was driven by a high abundance of *Wolbachia*. As *Wolbachia* is known to sometimes reach high titers in *D. sturtevanti*, this result is consistent with the literature (**Supplementary Figure 6**; Moreira, 2020). In the *bipectinata* complex, the enrichment in *Gammaproteobacteria* was mainly due to *Orbus hercynius* and *Orbaceae* sp. 3 (**Supplementary Figure 6**). Although these taxa are more commonly associated with native species, they were also detected at low abundance in other IAS pools and remain below the threshold defining core taxa, making their presence unsurprising.

In contrast, mycobiome class distribution was similarly dominated by *Saccharomycetes* and *Tremellomycetes* across IAS and native species. *Saccharomycetes* are well documented in *Drosophila* but *Tremellomycetes* remain comparatively understudied in this genus despite being reported in other insects, likely reflecting the limited number of studies on wild *Drosophila* mycobiomes (Chandler et al., 2012; Zhang et al., 2018; Cini et al., 2020; Cheng et al., 2023).

Overall, these results support differences in bacteriome composition and structure between IAS and native species and emphasise the need for finer taxonomic analyses to characterise these patterns better.

### Candidate taxa associated with native and invasive hosts

By examining abundance/prevalence, and global distribution patterns, we identified 38 bacterial taxa and 20 fungal taxa underlying community-level differences between native and IAS hosts.

Based on a literature review, we further excluded 11 taxa from this candidate list (see **Supplementary Table 8** for a summary). Some taxa, such as *Suttonella indologenes*, *Bartonella*, *Haemophilus*, *Erwinia*, and *Negativibacillus*, have environmental functions or ecologies incompatible with a true association with *Drosophila*, and most likeky correspond to environmental taxa (**Supplementary Table 8**). Others, including *Gammaproteobacteria*, *Burkholderiaceae*, *Tannerellaceae*, *Ruminococcaceae*, *Leotiomycetes* and *Saccharomycetales* were either assigned at too high a taxonomic rank or corresponded to highly diverse and/or poorly characterised groups, which prevents meaningful interpretation (**Supplementary Table 8**).

The remaining 47 taxa constitute our candidate list, they are summarised in **Table 3**. The list is composed of bacteria taxa : *Leuconostoc*, *Apibacter*, *Candidatus Soleaferrea*, *Cedecea*, *Orbus*, *Lacticaseibacillus*, *Komagataeibacter saccharivorans*, *Kozakia baliensis*, *Orbus sasakiae*, *Morganella morganii*, *Flavobacterium naphthae*, *Chishuiella*, *Spiroplasma*, *Candidatus Schmidhempelia bombi*, *Chryseobacterium*, *Enterococcus*, *Enterobacteriaceae*, *Orbaceae*, Cronobacter, *Orbus hercynius*, *Lactobacillus rossiae*, *Comamonadaceae*, *Pseudomonas*, *Dysgonomonas*, *Wolbachia*, *Acetobacter*, *Weeksellaceae*, *Commensalibacter* and *Commensalibacter intestini* (**Table 3**).

The candidate list is also composed of fungal groups: *Rhodotorula mucilaginosa*, *Candida californica*, *Diutina catenulata*, *Geotrichum*, *Candida gigantensis*, *Diutina bernalii*, *Gliocephalotrichum humicola*, *Hanseniaspora uvarum*, *Candida haemulonii*, *Pichia mandshurica*, *Saprochaete*, *Pichia bruneiensis*, *Clavispora fructus*, *Meyerozyma guiliermondii*, *Gliocephalotrichum mexicanum*, *Pichia sporocuriosa*, *Oligophagozyma castelli* and *Dipodascaceae* (**Table 3**).

These candidate taxa characterise microbiome differences between native and IAS hosts and provide a focused set of targets deserving future investigation, either in refining their taxonomic affiliation or because of a potential functional role associate them to their host.

### Limitations of this study and perspectives

As explained above, the respective effects of Host Species, Category, and Locality on bacteriome diversity, composition, and structure are difficult to disentangle. IAS and native species are necessarily represented by different host species, and most IAS individuals were collected in Cayenne, creating potential confounding between Category, Host Species, and Locality. Pairwise comparisons nevertheless supported independent effects of Host Species and Locality (Supplementary Tables 5 and 6), including significant locality differences after excluding Cayenne. The main uncertainty therefore concerns the Category effect. Several observations suggest that it may not be entirely explained by these confounding factors: some taxa were exclusively associated with IAS or differentially represented between IAS and native species despite the presence of native individuals in Cayenne, and no Category effect was detected for the mycobiome despite significant Host Species and Locality effects.

Nevertheless, our results cannot demonstrate a causal effect of invasion status independently of host identity and should instead be interpreted as consistent bacteriome differences between IAS and native species. This distinction is particularly important because Category was also phylogenetically structured: analysed IAS belonged to the melanogaster group, whereas native hosts belonged predominantly to the willistoni and saltans groups. Insufficient replication of the phylogenetically distinct IAS, *Z. indianus*, prevented us from reducing this confounding effect. A more balanced sampling design will therefore be necessary to disentangle invasion status, host identity, phylogeny, and locality.

Such a sampling design could also investigate potential microorganism exchange between native and IAS. Pathogens and parasites can be co-introduced with IAS and transmitted to native species (co-invasion), whereas IAS may acquire parasites or pathogens from native species, potentially affecting their establishment (spillback) (Taraschewski et al., 2006; Poulin et al., 2011; Hatcher et al., 2012; Lymbery et al., 2014). Because IAS occurred at low frequencies in some localities, future sampling could compare microbiomes of native hosts occurring with and without IAS to investigate such exchanges.

Our study was also restricted to adult flies and to bacterial and fungal communities.

Including larval stages and other microbial components would provide a more comprehensive view of microbiome dynamics. Viruses require substantially different methodological approaches and were therefore excluded from this study, while archaeal communities remain less well characterised than bacterial and fungal communities. Dedicated analyses of these components represent an important perspective for future studies.

Taxonomic and functional resolution also remain important limitations. Broad-range 16S rRNA and ITS markers do not provide uniform resolution across the considerable diversity of bacteria and fungi (Janda & Abbott, 2007; Schoch et al., 2012), and species boundaries can themselves be difficult to delineate (Taylor et al., 2000; Bobay, 2020). Incomplete reference databases further restrict species-level assignments (Locey & Lennon, 2016; Hawksworth & Lücking, 2017). Moreover, metabarcoding provides limited information on microbial gene content, expression, metabolic activity, or effects on host phenotype, preventing reliable functional inference from taxonomy alone.

Future analyses could improve taxonomic resolution using markers targeting candidate taxa, such as *dnaK*, *groEL*, and *rpoB* for *Acetobacter* (Li et al., 2014). Shotgun metagenomics and metagenome-assembled genomes (MAGs) could further improve genomic and functional resolution (Comeault et al., 2024), although their efficiency in host-associated samples is constrained by host DNA, sequencing depth, and microbial abundance, potentially limiting the recovery of rare taxa. Combining broad metabarcoding with targeted metagenomics or isolate-based characterization therefore represents a promising approach.

Ultimately, experimental validation, for example through controlled microbial inoculations, will be required to directly test the functional roles of candidate taxa, following characterization of microbiome differences between wild populations and laboratory isofemale lines.

## Conclusion

In this study, we showed that IAS of *Drosophila* are more abundant and diverse in anthropised localities such as Cayenne, while remaining scarce in rainforest environments. These patterns reveal a strong ecological contrast between native and invasive *Drosophila* across anthropised and rainforest localities, although the factors underlying this distribution remain unresolved.

By generating the first dataset from French Guiana of *Drosophila* diversity and their natural microbiomes, we identified reduced bacteriome diversity in IAS, along with marked differences in bacterial community composition and structure between host categories. In contrast, mycobiome differences were weaker at the community level but became apparent through finer-scale taxonomic analyses. Analysis of exclusive and core taxa revealed 47 bacterial and fungal candidate taxa that differed in their occurrence or core association between IAS and native hosts.

These candidates provide a basis for future functional investigations, but their roles remain to be experimentally tested. In particular, targeted taxonomic markers, long-read sequencing, metagenome-assembled genomes, and controlled microbial inoculation experiments will be needed to determine whether these taxa directly influence host adaptation, competitiveness, or invasion success.

Altogether, these findings establish a baseline characterization of microbiome variation between native and invasive *Drosophila* in French Guiana and provide candidate microbial taxa for future studies testing their ecological and functional significance.

## Acknowledgements

We thank Stephan Cassin for discussions and reflections on protocols, Guillaume Correà-Pimpao and Hugo Lorioux-Chevalier for field assistance, and Isabelle Clavereau, Francis Jiggins, and Marine Ginouves for laboratory assistance and advice.

We thank the Nouragues research field station (CNRS) which benefits from “Investissement d’Avenir” grants managed by Agence Nationale de la Recherche (AnaEE France ANR-24-INBS-0001; Labex CEBA ANR-10-LABX-25-01). Part of the specimens used in this study were collected in the Natural Reserve of Nouragues.

## Data Availability

The sequence datasets generated and/or analyzed during the current study are available in the EMBL Nucleotide Sequence Database (ENA), with Project accession PRJEB112398. https://www.ebi.ac.uk/ena/data/view/PRJEB112398 COI consensus (ERZ29525459) and OTU centroids sequences (ERZ29526552 and ERZ29525713) are available at http://www.ebi.ac.uk/ena/data/view/ERZ29526552 and http://www.ebi.ac.uk/ena/data/view/ERZ29525713.

All scripts used for data processing, statistical analyses, and figure generation are available at https://github.com/Thibault-Laffargue/Microbial-profiling-of-native-and-alien-Drosophila-in-French-Guiana. Other data types supporting the findings are available within the article and its supplementary files.

