## Supplementary data for "Microbial profiling of Drosophila in French Guiana reveals candidate microbial taxa associated with native and invasive host"

### Supplement

**Supplementary Table 1:** Primer list.

| Primer name | Primer sequence (5’ -> 3’) | Marker | References |
|---|---|---|---|
| COI F 106 | ATTCAGAATATCTATGTTCAG | *mt-COI* | Madi-Ravazzi et al. 2021 |
| COI R 154 | TTTAATTTTACCTGGATTTGG | *mt-COI* | Madi-Ravazzi et al. 2021 |
| BACT27F | AGAGTTTGATCMTGGCTCAG | 16s (V1 to V8) | Klindworth et al., 2013 |
| BACT1391R | GACGGGCGGTGWGTRCA | 16s (V1 to V8) | Klindworth et al., 2013 |
| OL4-F(ITS1F) | GCCTTGGTCATTTAGAGGAAGTAA | ITS1 to ITS2 | Gardes and Bruns, 1993; Chen et al., 2022 |
| OL4-R(ITS4KYO1) | TCCTCCGCTTWTTGWTWTGC | ITS1 to ITS2 | Chen et al., 2022 |

**Supplementary Table 2:** Table of COI primer and barcode.

| Name | **Barcode sequence** (5′ -> 3′) | **Full sequence** (5′ -> 3′) |
|---|---|---|
| oligo-DC-F1 | ATCAGTCAGACGC | ATCAGTCAGACGCATTCAGAATATCTATGTTCAG |
| oligo-DC-F2 | ACAACCATTGGCA | ACAACCATTGGCAATTCAGAATATCTATGTTCAG |
| oligo-DC-F3 | ACATCAGTAGTTC | ACATCAGTAGTTCATTCAGAATATCTATGTTCAG |
| oligo-DC-F4 | ACGCGCTCTTATA | ACGCGCTCTTATAATTCAGAATATCTATGTTCAG |
| oligo-DC-F5 | ACTTGTGCACCTG | ACTTGTGCACCTGATTCAGAATATCTATGTTCAG |
| oligo-DC-F6 | AGCTCGTGTCGTC | AGCTCGTGTCGTCATTCAGAATATCTATGTTCAG |
| oligo-DC-F7 | ACCTTCGCATAGA | ACCTTCGCATAGAATTCAGAATATCTATGTTCAG |
| oligo-DC-F8 | AGTGTGTAGCGCA | AGTGTGTAGCGCAATTCAGAATATCTATGTTCAG |
| oligo-DC-R1 | CAAGATCGGTACC | CAAGATCGGTACCTTTAATTTTACCTGGATTTGG |
| oligo-DC-R2 | GCAGTGCGAGTAG | GCAGTGCGAGTAGTTTAATTTTACCTGGATTTGG |
| oligo-DC-R3 | CAATGTTAATGGT | CAATGTTAATGGTTTTAATTTTACCTGGATTTGG |
| oligo-DC-R4 | CACCGGTAGAACC | CACCGGTAGAACCTTTAATTTTACCTGGATTTGG |
| oligo-DC-R5 | CAGAACAACGCAT | CAGAACAACGCATTTTAATTTTACCTGGATTTGG |
| oligo-DC-R6 | CATGCACACCTCT | CATGCACACCTCTTTTAATTTTACCTGGATTTGG |
| oligo-DC-R7 | CATCCTATCTCAT | CATCCTATCTCATTTTAATTTTACCTGGATTTGG |
| oligo-DC-R8 | GCGAGCGTCGAGA | GCGAGCGTCGAGATTTAATTTTACCTGGATTTGG |
| oligo-DC-R9 | CCAGCCTGACGGT | CCAGCCTGACGGTTTTAATTTTACCTGGATTTGG |
| oligo-DC-R10 | CCGAAGCCGGTTC | CCGAAGCCGGTTCTTTAATTTTACCTGGATTTGG |
| oligo-DC-R11 | CGTCGACATGATG | CGTCGACATGATGTTTAATTTTACCTGGATTTGG |
| oligo-DC-R12 | CCGTAGTGTTGAT | CCGTAGTGTTGATTTTAATTTTACCTGGATTTGG |

**Supplementary Table 3:** Table of 16S primer and barcode.

| Name | Barcode sequence (5′ -> 3′) | Full sequence (5′ -> 3′) |
|---|---|---|
| BACT27F_F1 | AACATGTGGTAAG | AACATGTGGTAAGAGAGTTTGATCMTGGCTCAG |
| BACT27F_F2 | ACAACCATTGGCA | ACAACCATTGGCAAGAGTTTGATCMTGGCTCAG |
| BACT27F_F3 | ACATCAGTAGTTC | ACATCAGTAGTTCAGAGTTTGATCMTGGCTCAG |
| BACT27F_F4 | ACGCGCTCTTATA | ACGCGCTCTTATAAGAGTTTGATCMTGGCTCAG |
| BACT27F_F5 | ACTTGTGCACCTG | ACTTGTGCACCTGAGAGTTTGATCMTGGCTCAG |
| BACT27F_F6 | AGACTCGATTGAG | AGACTCGATTGAGAGAGTTTGATCMTGGCTCAG |
| BACT27F_F7 | AGATTCTACACAA | AGATTCTACACAAAGAGTTTGATCMTGGCTCAG |
| BACT27F_F8 | AGCCTAGCCAACT | AGCCTAGCCAACTAGAGTTTGATCMTGGCTCAG |
| BACT1391R_R1 | CAAGATCGGTACC | CAAGATCGGTACCGACGGGCGGTGWGTRCA |
| BACT1391R_R2 | CAATCCTCAAGAG | CAATCCTCAAGAGGACGGGCGGTGWGTRCA |
| BACT1391R_R3 | CAATGTTAATGGT | CAATGTTAATGGTGACGGGCGGTGWGTRCA |
| BACT1391R_R4 | CACCGGTAGAACC | CACCGGTAGAACCGACGGGCGGTGWGTRCA |
| BACT1391R_R5 | CAGAACAACGCAT | CAGAACAACGCATGACGGGCGGTGWGTRCA |
| BACT1391R_R6 | CATGCACACCTCT | CATGCACACCTCTGACGGGCGGTGWGTRCA |
| BACT1391R_R7 | CATGCCTTGATAC | CATGCCTTGATACGACGGGCGGTGWGTRCA |
| BACT1391R_R8 | CATTATATAGCCA | CATTATATAGCCAGACGGGCGGTGWGTRCA |
| BACT1391R_R9 | CCAGCCTGACGGT | CCAGCCTGACGGTGACGGGCGGTGWGTRCA |
| BACT1391R_R10 | CCGAAGCCGGTTC | CCGAAGCCGGTTCGACGGGCGGTGWGTRCA |
| BACT1391R_R11 | CCGAGCAGCTGTT | CCGAGCAGCTGTTGACGGGCGGTGWGTRCA |
| BACT1391R_R12 | CCGTAGTGTTGAT | CCGTAGTGTTGATGACGGGCGGTGWGTRCA |

**Supplementary Table 4:** Table of ITS primer and barcode.

| Name | Barcode sequence (5′ -> 3′) | **Full sequence** (5′ -> 3′) |
|---|---|---|
| IFungITS_F1 | AACATGTGGTAAG | AACATGTGGTAAGGCCTTGGTCATTTAGAGGAAGTAA |
| IFungITS_F2 | ACAACCATTGGCA | ACAACCATTGGCAGCCTTGGTCATTTAGAGGAAGTAA |
| IFungITS_F3 | ACATCAGTAGTTC | ACATCAGTAGTTCGCCTTGGTCATTTAGAGGAAGTAA |
| IFungITS_F4 | ACGCGCTCTTATA | ACGCGCTCTTATAGCCTTGGTCATTTAGAGGAAGTAA |
| IFungITS_F5 | ACTTGTGCACCTG | ACTTGTGCACCTGGCCTTGGTCATTTAGAGGAAGTAA |
| IFungITS_F6 | AGACTCGATTGAG | AGACTCGATTGAGGCCTTGGTCATTTAGAGGAAGTAA |
| IFungITS_F7 | AGATTCTACACAA | AGATTCTACACAAGCCTTGGTCATTTAGAGGAAGTAA |
| IFungITS_F8 | AGCCTAGCCAACT | AGCCTAGCCAACTGCCTTGGTCATTTAGAGGAAGTAA |
| IFungITS_R1 | CAAGATCGGTACC | CAAGATCGGTACCTCCTCCGCTTWTTGWTWTGC |
| IFungITS_R2 | CAATCCTCAAGAG | CAATCCTCAAGAGTCCTCCGCTTWTTGWTWTGC |
| IFungITS_R3 | CAATGTTAATGGT | CAATGTTAATGGTTCCTCCGCTTWTTGWTWTGC |
| IFungITS_R4 | CACCGGTAGAACC | CACCGGTAGAACCTCCTCCGCTTWTTGWTWTGC |
| IFungITS_R5 | CAGAACAACGCAT | CAGAACAACGCATTCCTCCGCTTWTTGWTWTGC |
| IFungITS_R6 | CATGCACACCTCT | CATGCACACCTCTTCCTCCGCTTWTTGWTWTGC |
| IFungITS_R7 | CATGCCTTGATAC | CATGCCTTGATACTCCTCCGCTTWTTGWTWTGC |
| IFungITS_R8 | CATTATATAGCCA | CATTATATAGCCATCCTCCGCTTWTTGWTWTGC |
| IFungITS_R9 | CCAGCCTGACGGT | CCAGCCTGACGGTTCCTCCGCTTWTTGWTWTGC |
| IFungITS_R10 | CCGAAGCCGGTTC | CCGAAGCCGGTTCTCCTCCGCTTWTTGWTWTGC |
| IFungITS_R11 | CCGAGCAGCTGTT | CCGAGCAGCTGTTTCCTCCGCTTWTTGWTWTGC |
| IFungITS_R12 | CCGTAGTGTTGAT | CCGTAGTGTTGATTCCTCCGCTTWTTGWTWTGC |

**Supplementary Table 5:** Significant Kruskal-Wallis and Wilcoxon pairwise results for observed specific richness and Shannon index.

| Dataset | Model | Test | Term | P-value |
|---|---|---|---|---|
| bacteriome - global | Observed ~ HostSpecies | Kruskal-Wallis | HostSpecies | 4.29E-05 |
| | | Wilcoxon pairwise | pau VS Bip | 0.001767 |
| | | | stu VS pau | 0.003178 |
| | Observed ~ Category | Kruskal-Wallis | Category | 0.010387 |
| | Observed ~ Locality | Kruskal-Wallis | Locality | 0.000543 |
| | | Wilcoxon pairwise | Cayenne VS Bélizon | 0.005391 |
| | | | Nouragues VS Bélizon | 0.006501 |
| | Shannon ~ HostSpecies | Kruskal-Wallis | HostSpecies | 6.78E-06 |
| | | Wilcoxon pairwise | pau VS Bip | 2.84E-06 |
| | | | stu VS pau | 0.004924 |
| | | | Wil VS Bip | 0.023529 |
| | Shannon ~ Category | Kruskal-Wallis | Category | 7.82E-06 |
| | | Wilcoxon pairwise | Native VS Invasive | 9.79E-07 |
| | Shannon ~ Locality | Kruskal-Wallis | Locality | 0.000346 |
| | | Wilcoxon pairwise | Cayenne VS Bélizon | 0.000557 |
| | | | Kaw VS Cayenne | 0.003243 |
| | | | Nouragues VS Cayenne | 0.004269 |
| | Pielou ~ HostSpecies | Kruskal-Wallis | HostSpecies | 1.63E-05 |
| | | Wilcoxon pairwise | pau VS Bip | 7.16E-06 |
| | | | stu VS pau | 0.020456 |
| | | | Wil VS Bip | 0.040336 |
| | | | Wil VS stu | 0.095238 |
| | Pielou ~ Category | Kruskal-Wallis | Category | 8.51E-06 |
| | | Wilcoxon pairwise | Native VS Invasive | 1.12E-06 |
| | Pielou ~ Locality | Kruskal-Wallis | Locality | 0.000455 |
| | | Wilcoxon pairwise | Cayenne VS Bélizon | 0.000902 |
| | | | Kaw VS Cayenne | 0.003243 |
| | | | Nouragues VS Cayenne | 0.004269 |
| mycobiome - global | Observed ~ Locality | Kruskal-Wallis | Locality | 0.00112 |
| | | Wilcoxon pairwise | Nouragues VS Bélizon | 0.001624 |
| | Shannon ~ Locality | Kruskal-Wallis | Locality | 0.005896 |
| | | Wilcoxon pairwise | Kaw VS Bélizon | 0.010016 |
| | | | Nouragues VS Bélizon | 0.008462 |
| | Pielou ~ Locality | Kruskal-Wallis | Locality | 0.043054 |

**Supplementary Table 6:** Significant Permanova results for ordination analyses.

| Dataset | Model | Test | Term | P-value | R2 |
|---|---|---|---|---|---|
| bacteriome | UniFrac | multivariate analogue of levene's test | HostSpecies | 4.8E-08 | |
| | | ~ HostSpecies X Locality | HostSpecies | 0.001 | 0.1551 |
| | | | Locality | 0.001 | 0.2747 |
| | | | Locality:HostSpecies | 0.031 | 0.0757 |
| | | ~ Locality X Category | Category | 0.001 | 0.0443 |
| | | | Locality | 0.001 | 0.2747 |
| | | ~ Category X Locality | Category | 0.001 | 0.2476 |
| | | | Locality | 0.005 | 0.0714 |
| | | ~ HostSpecies | Bip vs pau | 0.006 | 0.3245 |
| | | | Bip vs stu | 0.006 | 0.2902 |
| | | | Bip vs Wil | 0.006 | 0.3362 |
| | | | pau vs stu | 0.006 | 0.1388 |
| | | | pau vs Wil | 0.018 | 0.0547 |
| | | | Wil vs stu | 0.036 | 0.2578 |
| | | ~ Locality | Bélizon vs Nouragues | 0.012 | 0.0801 |
| | | | Cayenne vs Bélizon | 0.006 | 0.305 |
| | | | Cayenne vs Kaw | 0.006 | 0.3323 |
| | | | Cayenne vs Nouragues | 0.006 | 0.2525 |
| | Weighted UniFrac | multivariate analogue of levene's test | HostSpecies | 0.04929 | |
| | | ~ HostSpecies X Locality | HostSpecies | 0.001 | 0.1935 |
| | | | Locality | 0.001 | 0.4496 |
| | | | Locality:HostSpecies | 0.026 | 0.0631 |
| | | ~ Locality X Category | Category | 0.001 | 0.1059 |
| | | | Locality | 0.001 | 0.4496 |
| | | ~ Category X Locality | Category | 0.001 | 0.5334 |
| | | ~ HostSpecies | Bip vs pau | 0.006 | 0.6575 |
| | | | Bip vs stu | 0.012 | 0.3573 |
| | | | Bip vs Wil | 0.006 | 0.5724 |
| | | | pau vs stu | 0.006 | 0.2277 |
| | | ~ Locality | Cayenne vs Bélizon | 0.006 | 0.574 |
| | | | Cayenne vs Kaw | 0.006 | 0.515 |
| | | | Cayenne vs Nouragues | 0.012 | 0.3325 |
| bacteriome native only | UniFrac | ~ HostSpecies X Locality | HostSpecies | 0.001 | 0.1554 |
| | | | Locality | 0.002 | 0.1155 |
| | | | Locality:HostSpecies | 0.022 | 0.1043 |
| | Weighted UniFrac | | HostSpecies | 0.001 | 0.2321 |
| | | | Locality:HostSpecies | 0.037 | 0.1563 |

| Dataset | Model | Test | Term | P-value | R2 |
|---|---|---|---|---|---|
| mycobiome | Bray-Curtis | multivariate analogue of levene's test | HostSpecies | 0.0168 | |
| | | | Locality | 0.00655 | |
| | | ~ HostSpecies X Locality | HostSpecies | 0.001 | 0.16007 |
| | | | Locality | 0.001 | 0.16672 |
| | | ~ Locality X Category | Locality | 0.001 | 0.16672 |
| | | ~ HostSpecies | Bip vs pau | 0.03 | 0.08435 |
| | | | Bip vs stu | 0.045 | 0.24032 |
| | | | pau vs stu | 0.075 | 0.07012 |
| | | | pau vs Sal | 0.015 | 0.11184 |
| | | ~ Locality | Bélizon vs Nouragues | 0.006 | 0.09951 |
| | | | Cayenne vs Bélizon | 0.006 | 0.08823 |
| | | | Cayenne vs Kaw | 0.006 | 0.21169 |
| | | | Cayenne vs Nouragues | 0.042 | 0.12924 |
| | | | Kaw vs Bélizon | 0.012 | 0.10803 |
| | Jaccard | multivariate analogue of levene's test | Category | 0.02594 | |
| | | | HostSpecies | 2.9E-06 | |
| | | ~ HostSpecies X Locality | HostSpecies | 0.001 | 0.13339 |
| | | | Locality | 0.001 | 0.12981 |
| | | | Locality:HostSpecies | 0.043 | 0.10421 |
| | | ~ Locality X Category | Category | 0.142 | 0.02257 |
| | | | Locality | 0.001 | 0.12981 |
| | | | Locality:Category | 0.001 | 0.0361 |
| | | ~ HostSpecies | Bip vs pau | 0.015 | 0.05245 |
| | | | pau vs stu | 0.015 | 0.06436 |
| | | | pau vs Sal | 0.09 | 0.06011 |
| | | ~ Locality | Bélizon vs Nouragues | 0.006 | 0.10361 |
| | | | Cayenne vs Bélizon | 0.006 | 0.08338 |
| | | | Cayenne vs Kaw | 0.072 | 0.08121 |
| | | | Cayenne vs Nouragues | 0.03 | 0.09826 |
| | | | Kaw vs Bélizon | 0.006 | 0.06607 |
| | | | Kaw vs Nouragues | 0.018 | 0.09965 |

**Supplementary Table 7:** List of taxa found exclusively in natives and invasive respectively.

| Exclusive to | Dataset | Species | Order | Abundance max | Abundance min | Prevalence | Host_species |
|---|---|---|---|---|---|---|---|
| Native | Bacteriome | Orbus sasakiae | Enterobacterales | 0.00136 | 0.0001338 | 0.6363636 | pau; Wil |
| | | Candidatus Soleaferrea sp. 1 | Oscillospirales | 0.010492 | 6.843E-05 | 0.5227273 | pau ; stu; Wil |
| | | Burkholderiaceae sp. 1 | Burkholderiales | 0.019956 | 6.196E-05 | 0.3636364 | pau; Wil |
| | | Orbus sp. 3 | Enterobacterales | 0.001425 | 0.0001724 | 0.3409091 | pau; Wil |
| | | Orbus sp. 2 | Enterobacterales | 0.008434 | 6.196E-05 | 0.3181818 | pau; Wil |
| | | Kozakia baliensis | Acetobacterales | 0.024929 | 0.0001338 | 0.2272727 | pau ; stu; Wil |
| | | Morganella morganii | Enterobacterales | 0.01192 | 6.196E-05 | 0.2272727 | pau ; stu |
| | | Apibacter sp. 1 | Flavobacteriales | 0.011667 | 0.0001976 | 0.2045455 | pau ; stu; Wil |
| | | Lacticaseibacillus sp. 1 | Lactobacillales | 0.001881 | 7.095E-05 | 0.2045455 | pau |
| | | Lacticaseibacillus sp. 2 | Lactobacillales | 0.001381 | 6.843E-05 | 0.2045455 | pau |
| | | Orbus sp. 1 | Enterobacterales | 0.001406 | 6.843E-05 | 0.1818182 | pau |
| | | Komagataeibacter saccharivorans | Acetobacterales | 0.025998 | 0.0003367 | 0.1590909 | pau ; stu |
| | | Cedecea sp. 1 | Enterobacterales | 0.000754 | 6.196E-05 | 0.1590909 | pau |
| | | Candidatus Soleaferrea sp. 2 | Oscillospirales | 0.00053 | 6.196E-05 | 0.1590909 | pau; Wil |
| | | Bartonella sp. 1 | Hyphomicrobiales | 0.003017 | 0.000369 | 0.1363636 | pau ; stu |
| | | Tannerellaceae sp. 1 | Bacteroidales | 0.012579 | 0.0005271 | 0.1136364 | pau |
| | | Haemophilus sp. 1 | Enterobacterales | 0.001954 | 0.0001369 | 0.1136364 | pau ; stu |
| | | Gammaproteobacteria sp. 1 | Gammaproteobacteria sp. | 0.004885 | 0.0004042 | 0.0909091 | pau ; stu; Wil |
| | | Flavobacterium naphthae | Flavobacteriales | 0.008929 | 0.0021186 | 0.0454545 | pau |
| | Mycobiome | Leotiomycetes sp. 1 | Leotiomycetes sp. | 0.121807 | 0.0003591 | 0.3414634 | pau; Wil |
| | | Gliocephalotrichum humicola | Hypocreales | 0.027923 | 0.000388 | 0.2195122 | pau |
| | | Candida_californica | Saccharomycetales | 0.004032 | 0.0004822 | 0.2195122 | pau ; stu |
| | | Geotrichum sp. | Saccharomycetales | 0.039487 | 0.0001138 | 0.1219512 | pau |
| | | Diutina bernalii | Saccharomycetales | 0.084195 | 0.0002313 | 0.097561 | pau |
| | | Candida gigantensis | Saccharomycetales | 0.027332 | 0.0005084 | 0.097561 | pau |
| | | Rhodotorula mucilaginosa | Sporidiobolales | 0.005455 | 0.0003591 | 0.097561 | pau |
| | | Diutina catenulata | Saccharomycetales | 0.019197 | 0.01576 | 0.0487805 | pau |
| Invasive | Bacteriome | Suttonella indologenes | Cardiobacteriales | 0.000435 | 7.151E-05 | 0.5714286 | Bip |
| | | Leuconostoc sp. 1 | Lactobacillales | 0.000523 | 4.53E-05 | 0.5 | Bip |

**Supplementary Table 8:** taxa excluded from the candidate list based on bibliography.

| Kingdom | Criteria | Taxa | Sources | Exclusion reason | Microbe atlas distribution |
|---|---|---|---|---|---|
| Bacteria | IAS exclusive | *Suttonella indologenes* | Sevencan et al., 2018 | acquisition in anthropized habitats | Commonly found in latin america |
| | Native exclusive | *Gammaproteobacteria* | Coenye, 2014; Ormerod et al., 2016; Mazzucco and Schlötterer, 2021 | Family rank is to high limiting functional interpretation | Cosmopolitan |
| | | *Burkholderiaceae* | | | |
| | | *Tannerellaceae* | | | |
| | | *Bartonella* | and Dehio, 2012; Dingle and Clarridge, 2014 | Environmental | Cosmopolitan |
| | | *Haemophilus* | | | |
| | Core native but not core IAS | *Erwinia* | 2011; Valles et al., 2021; Staubach et al., 2013 | Environmental or transient | Cosmopolitan |
| | | *Negativibacillus* | | | |
| | | *Ruminococcaceae* | | Family rank is to high limiting functional interpretation | Not documented |
| Fungi | Native exclusive | *Leotiomycetes* | Osono, 2006; Webster and Weber, 2012 | Class rank is to high limiting functional interpretation | Cosmopolitan |
| | Core native but not core IAS | *Meyerozyma guilliermondii* | Chandler et al., 2012; Stamps et al., 2012 | complex, inconsistent with our results | Cosmopolitan |
| | | *Saccharomycetales* | | Class rank is to high limiting functional interpretation | Cosmopolitan |

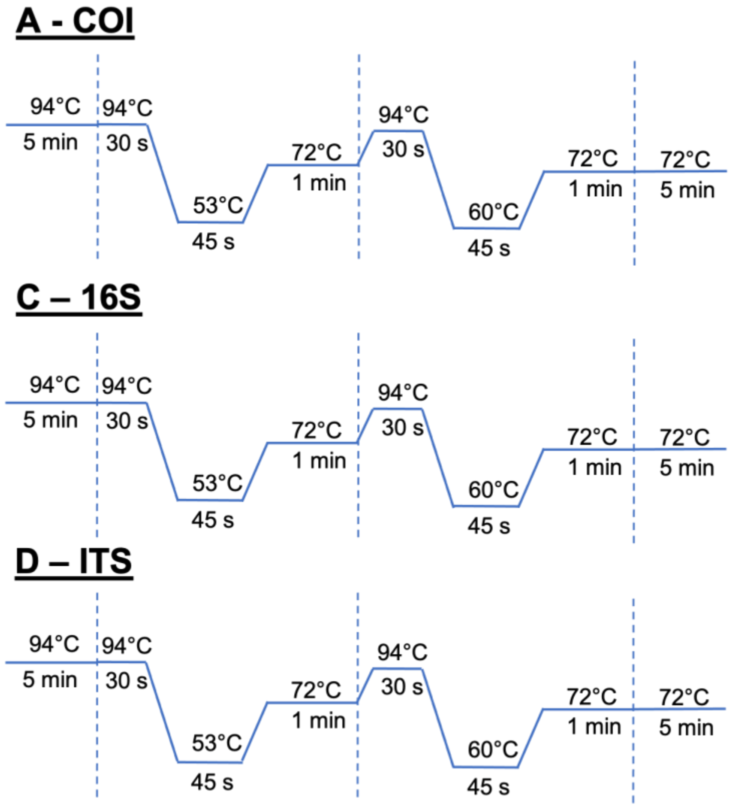


**B - COI**

| Components | Master Mix for 1 samples |
|---|---|
| H20 | 3.7 µL |
| 2X Euromedex | 10 µL |
| MgCl2 25mM | 1.2 µL |
| Primer F 5uM | 2 µL |
| Primer R 5uM | 2 µL |
| Template | 1.1 µL |
| Total volume | 20 µL |

**E – 16s and ITS**

| Component | Master Mix for 1 samples |
|---|---|
| 10X Amplus buffer | 2.5 µL |
| dNTP mix 5mM | 2.5 µL |
| Amplus DNA pol | 0.2 µL |
| Qsp water 25uL | 6.8 µL |
| 40ng DNA | 10 µL |
| Primer F 5uM | 1.5 µL |
| Primer R 5uM | 1.5 µL |
| Total volume | 25 µL |

**Supplementary Figure 1:** PCR condition.

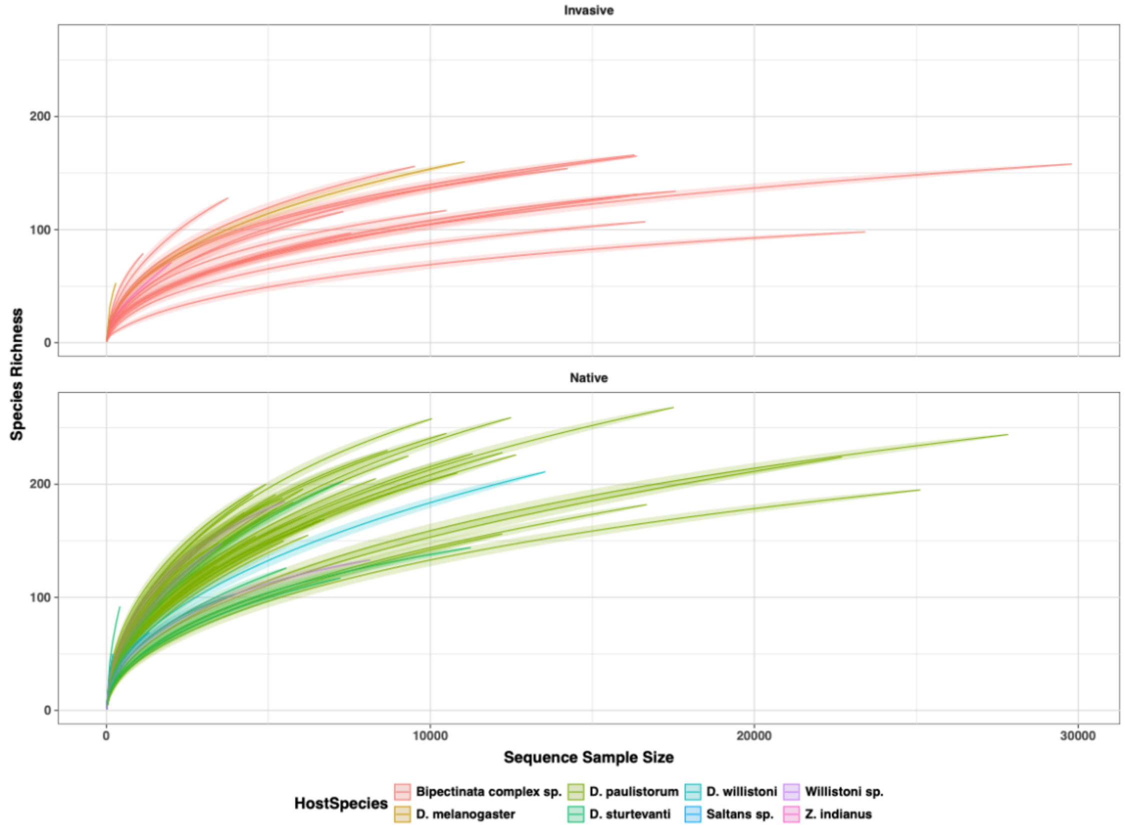


**Supplementary Figure 2:** 16S Rarefaction curve.

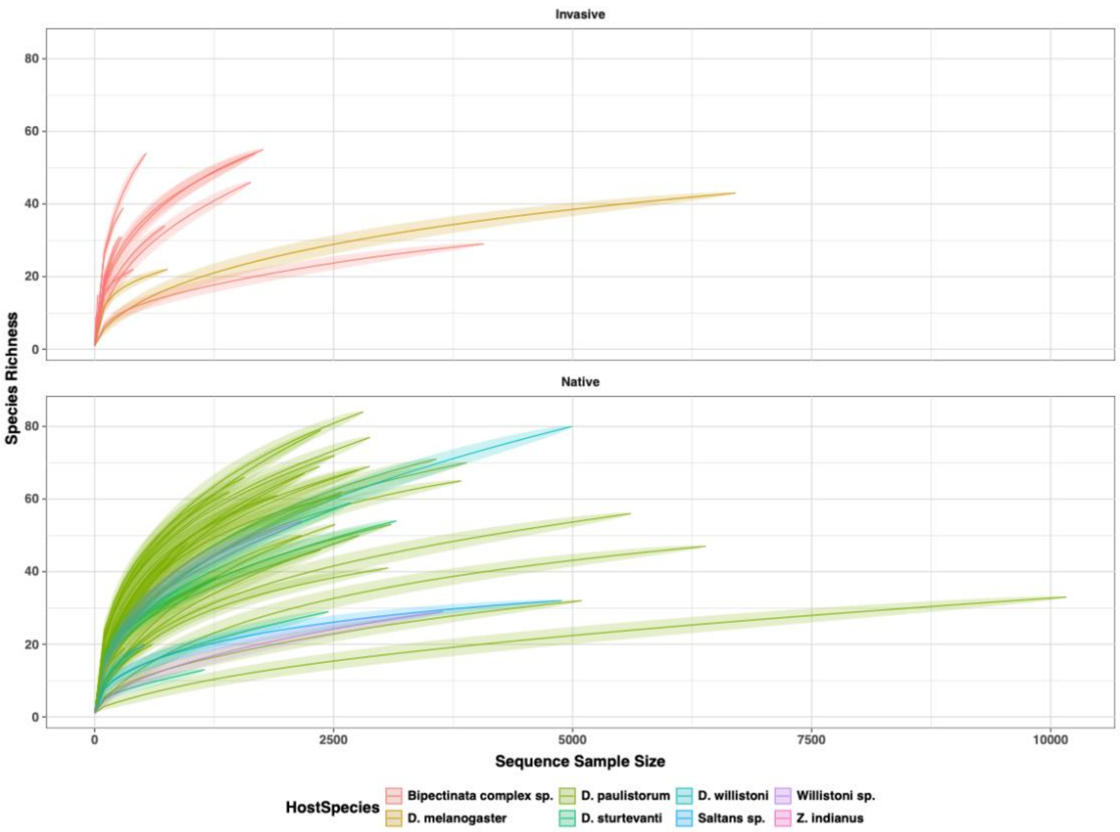


**Supplementary Figure 3:** ITS Rarefaction curve.

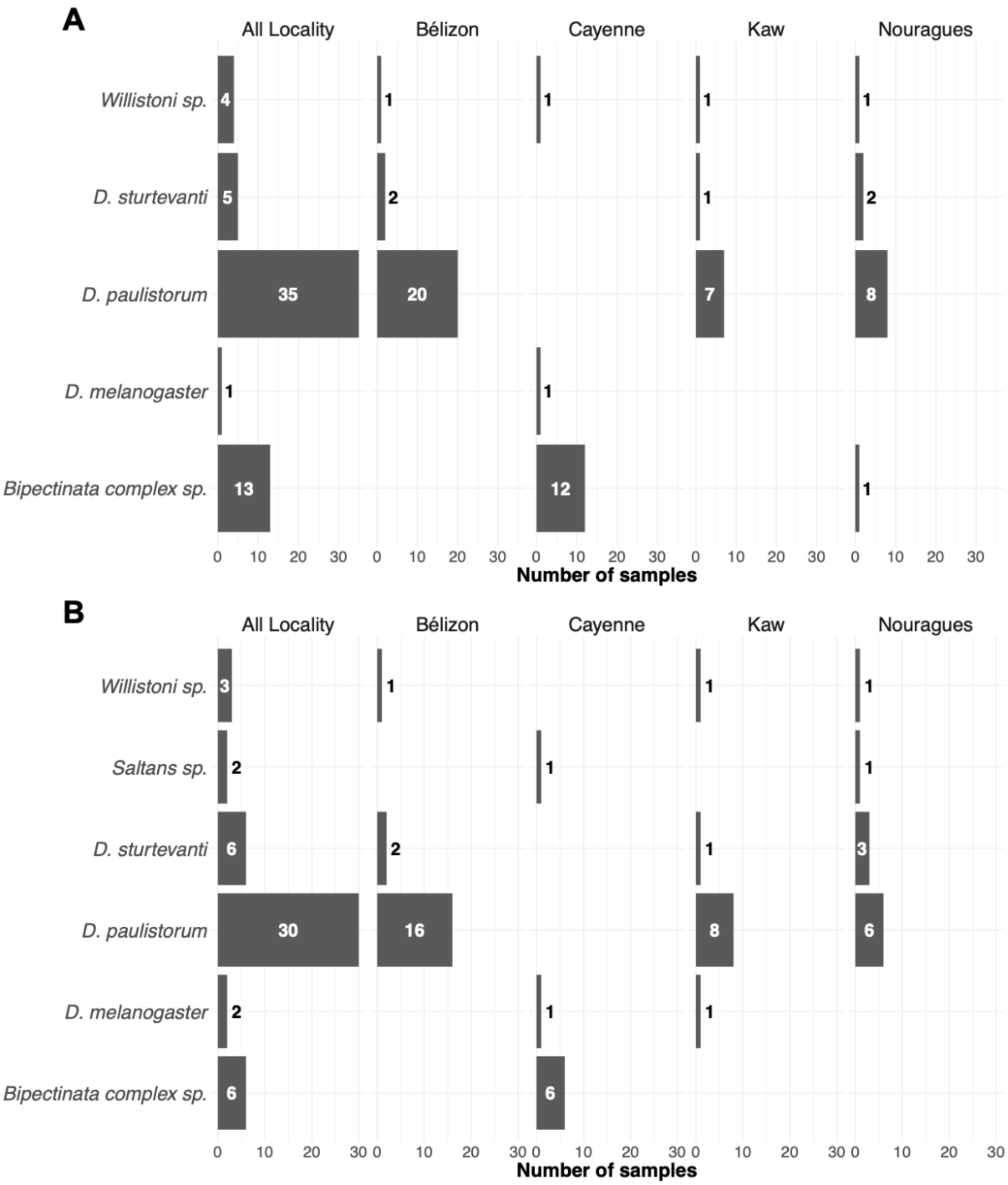


**Supplementary Figure 4:** number of pools, by locality and host species (A: 16s; B: ITS).

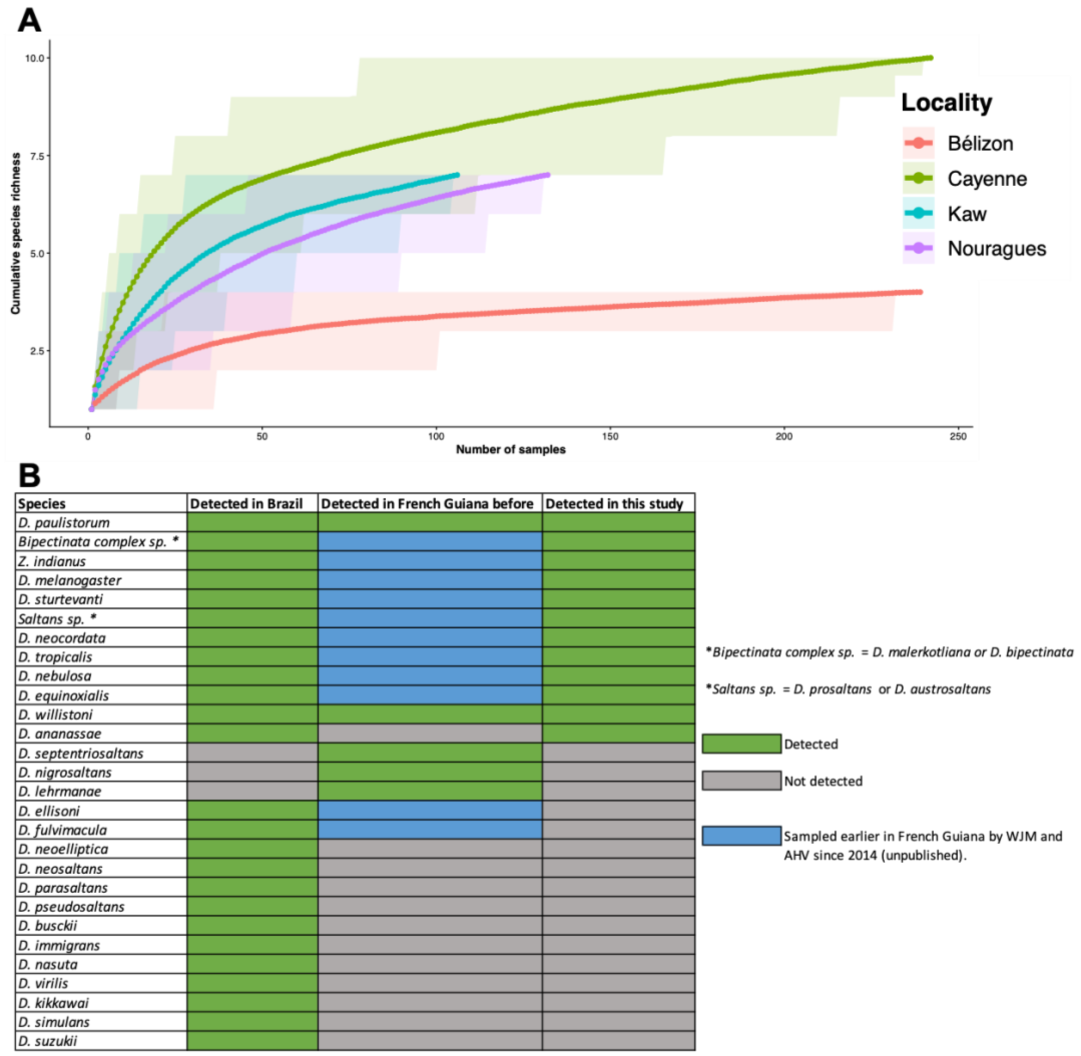


**Supplementary Figure 5: Evaluation of Drosophila sampling completeness. (A)** Species accumulation curves for each locality, showing species richness as a function of the cumulative number of samples. Curves were generated using 1,000 random permutations of sample order, and each point represents the mean cumulative species richness across permutations. Shaded areas represent the 95% permutation intervals. The curves approach saturation at all localities, suggesting that a substantial proportion of the species detectable with our sampling method was captured. **(B)** Comparison between Drosophila species detected in the present study and species previously reported from French Guiana and Brazil, a neighboring country. Previous occurrence records were compiled from Yuzuki and Tidon (2020), Baião et al. (2023), Prediger et al. (2024), and Papachristos et al. (2025), Montoliu-Nerin et al. (2026) and https://evolgen.biol.se.tmu.ac.jp/DrosWLD/. In blue are indicated species collected and *CO1*-barcoded by Wolfgang J. Miller and Aurélie Hua-Van in their systematic sampling trips in French Guiana since 2014 (unpublished). These studies were, however, not comprehensive surveys of native and invasive Drosophila communities but only reported as collected in French Guiana or Brazil. Species frequently detected in previous studies were also detected here. Our sampling did not detect five species previously described in French Guiana, which are considered to be infrequently collected or were collected in Saül, an inland locality geographically distant from the sites sampled here. We also identified one species that had never been described in French Guiana. Differences between previous records and the present study may reflect differences in sampling locations and methods as different baits were used. Many species detected in our study had previously been reported from Brazil, while numerous species reported from Brazil were not detected. Given the large geographic extent of Brazil and its diversity of biomes, Brazilian records should therefore be interpreted as evidence of regional occurrence rather than as an expectation of species presence in French Guiana.

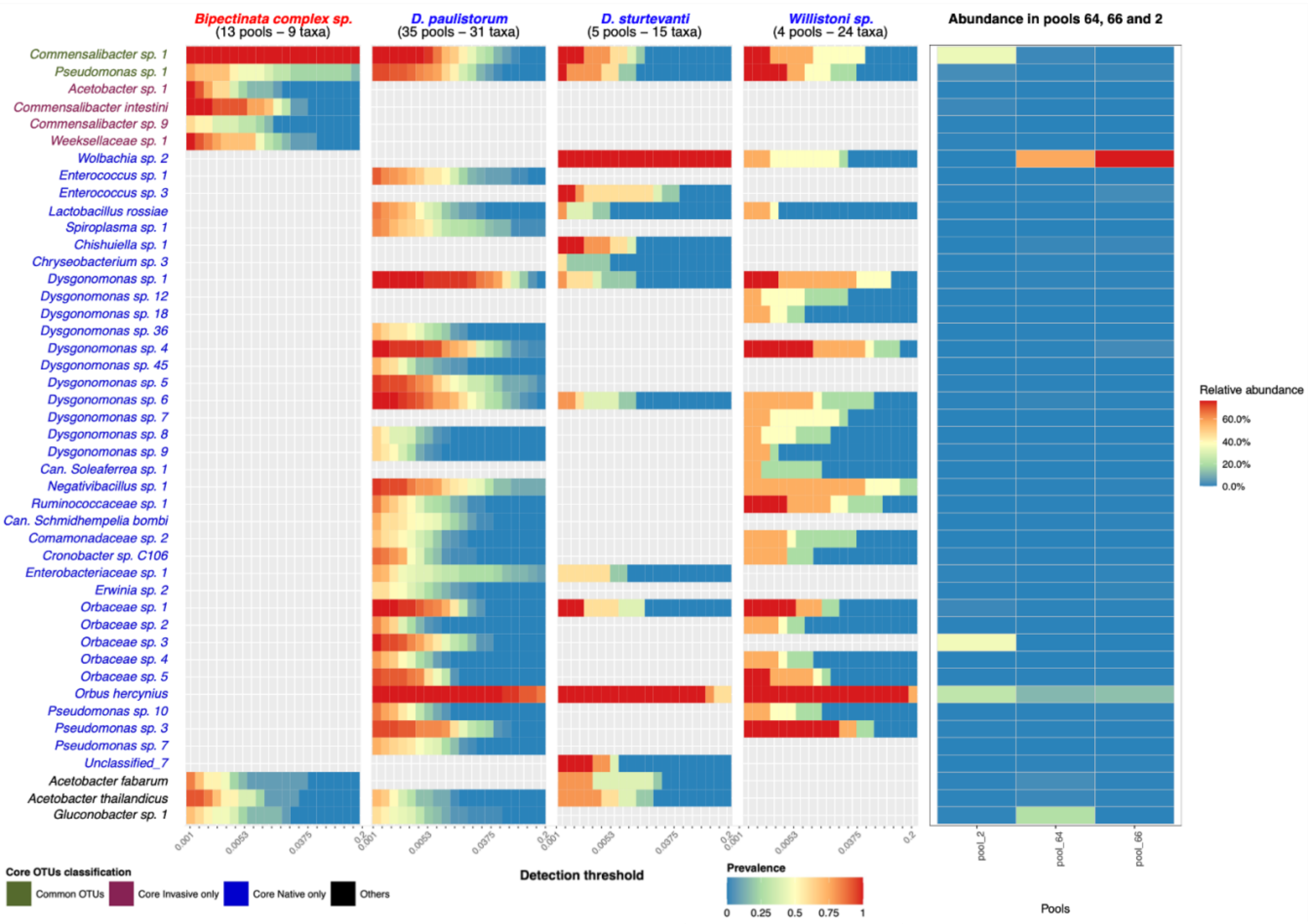


**Supplementary Figure 6:** unsimilar pool composition comparison. Pool 2 is a Bipectinata pool from Cayenne whose bacteria class composition differs from other IAS. Pool 64 and 66 are *D. sturtevanti* pools from Nouragues whose bacteria class composition differs from other native.
